# Stage-specific CAR-mediated signaling generates naïve-like, TCR-null CAR T cells from induced pluripotent stem cells

**DOI:** 10.1101/2024.11.25.624041

**Authors:** Sang Pil Yoo, Xuegang Yuan, Claire Engstrom, Patrick Chang, Suwen Li, Lindsay Lathrop, Jessica Lagosh, Satvik Saripalli, Anthony Azzun, Vincenzo Calvanese, Hanna Mikkola, Christopher Seet, Donald B. Kohn, Gay M. Crooks

## Abstract

Genetically modified, induced pluripotent stem cells (iPSCs) offer a promising allogeneic source for the generation of functionally enhanced, chimeric antigen receptor (CAR) T cells. However, the signaling of CARs during early T cell development and the removal of the endogenous T cell receptor required to prevent alloreactivity pose significant challenges to the production of mature conventional CAR T cells from iPSCs. Here, we show that TCR-null, CD8αβ CAR T cells can be efficiently generated from iPSCs by engineering stage-specific onset of CAR expression and signaling to both permit conventional T cell development and to induce efficient positive selection. CAR T cells produced using this approach displayed a uniform, naïve T cell phenotype and demonstrated superior antigen-specific cytotoxicity compared to iPSC-derived effector memory CAR T cells. Multimodal sequencing revealed CAR-mediated positive selection induced the persistent upregulation of key transcription factors involved in naïve T cell development. Achieving precise control of CAR expression and signaling in developmentally sensitive T precursors will be critical to realizing the full potential for “off-the-shelf”, iPSC-derived cellular therapies.

## Introduction

Genetically engineered, induced pluripotent stem cells (iPSCs) are under active investigation as an off-the shelf, allogeneic source of therapeutic T cells. The unique self-renewing capacity of iPSCs enables precise editing of multiple genes to enhance function and overcome critical immune barriers of an allogenic product^1,2^. However, the introduction or perturbation of biologically active genes can pose challenges to recapitulating conventional T cell differentiation *in vitro* from iPSCs. Inappropriate signaling or transcriptional cues provided at critical stages of development can result in a block in differentiation or diversion to an undesired lineage^3–5^.

These challenges have been reported in early attempts to generate chimeric antigen receptor (CAR) T cells from iPSCs. Unlike CAR T products derived from mature, peripheral blood T cells, the constitutive expression of a CAR during iPSC differentiation can divert development from the conventional T lineage and into innate pathways, resulting in the generation of CAR-expressing CD8αɑ, γδ T, type 2 innate lymphoid cells (ILC2s), and natural killer (NK) cells^6–9^. This diversion from conventional T cell differentiation has been attributed to signaling of certain CARs (particular those with high levels of antigen-independent or tonic signaling) in developmentally sensitive precursors^10^.

Solutions proposed to address the CAR-mediated block during conventional T differentiation have largely focused on altering the expression and design of CARs. We and others have shown that lowering the level of CAR expression during the early stages of T cell differentiation can reduce the degree of lineage diversion^7,11^. Alternatively, attenuating CAR signaling by replacing the CD28 co-stimulatory domain with the 4-1BB domain or selectively mutating motifs presented on the CD3ζ domain have been shown to allow some conventional T development^7,9,12^. However, modifying the CAR design and signaling may inadvertently and unpredictably come at the cost of reduced CAR T cytotoxicity, persistence, and long-term tumor control^13–15^. As a result, these strategies require rigorous and iterative empirical studies of each proposed CAR design within the context of the differentiation system used.

Complicating the production of iPSC-derived CAR T cells further, additional genetic edits necessary to address alloreactivity and allo-rejection disrupt the normal cellular processes required for T cell maturation. Specifically, preventing the expression of endogenous T cell receptors (TCRs) and major histocompatibility complexes (MHCs) removes the molecular machinery necessary for T cell precursors to achieve positive selection and complete development^16–18^. These challenges prompt the need for gene engineering approaches that carefully consider the exact timing, strength, and type of signals required at each stage of differentiation.

We report a novel gene editing approach that simultaneously addresses both CAR-mediated fate diversion and arrest in T cell maturation in the absence of TCR expression, that is compatible with a diverse range of CAR designs and modifications. Our approach leveraged the tightly regulated expression of endogenous genes to precisely control the onset of CAR expression at distinct stages of T cell development. Using this approach, innate fate diversion was prevented by restricting CAR expression to T cells and their precursors after commitment to the conventional developmental pathway. To rescue development in the absence of endogenous TCRs, CAR-mediated positive selection signals were engineered by introducing the CAR cognate antigen in the accessory cells of the artificial thymic organoid (ATO) system. By combining these strategies, we show the robust generation of TCRαβ-CD3- CD8αβ+ CAR T cells that possess a uniform, naïve-like phenotype and robust antigen-specific cytotoxicity. In summary, we outline strategies to overcome the biologic barriers to generating iPSC-derived, non-alloreactive CAR T cells, broadening the range of tumor targets and diseases that can be addressed by stem cell therapies.

## Results

### Establishing stage-specific transgene expression during T cell differentiation from iPSCs

To delay the onset of CAR expression during iPSC differentiation, we used CRISPR/Cas9-mediated homology directed repair (HDR) to insert transgenes into the 3’ untranslated region (UTR) of chosen endogenous genes that exhibit optimal stage-specific expression during T cell development; thus, transgene expression was co-regulated with that of the endogenous gene **(Fig. 1a).**

**Figure 1:**
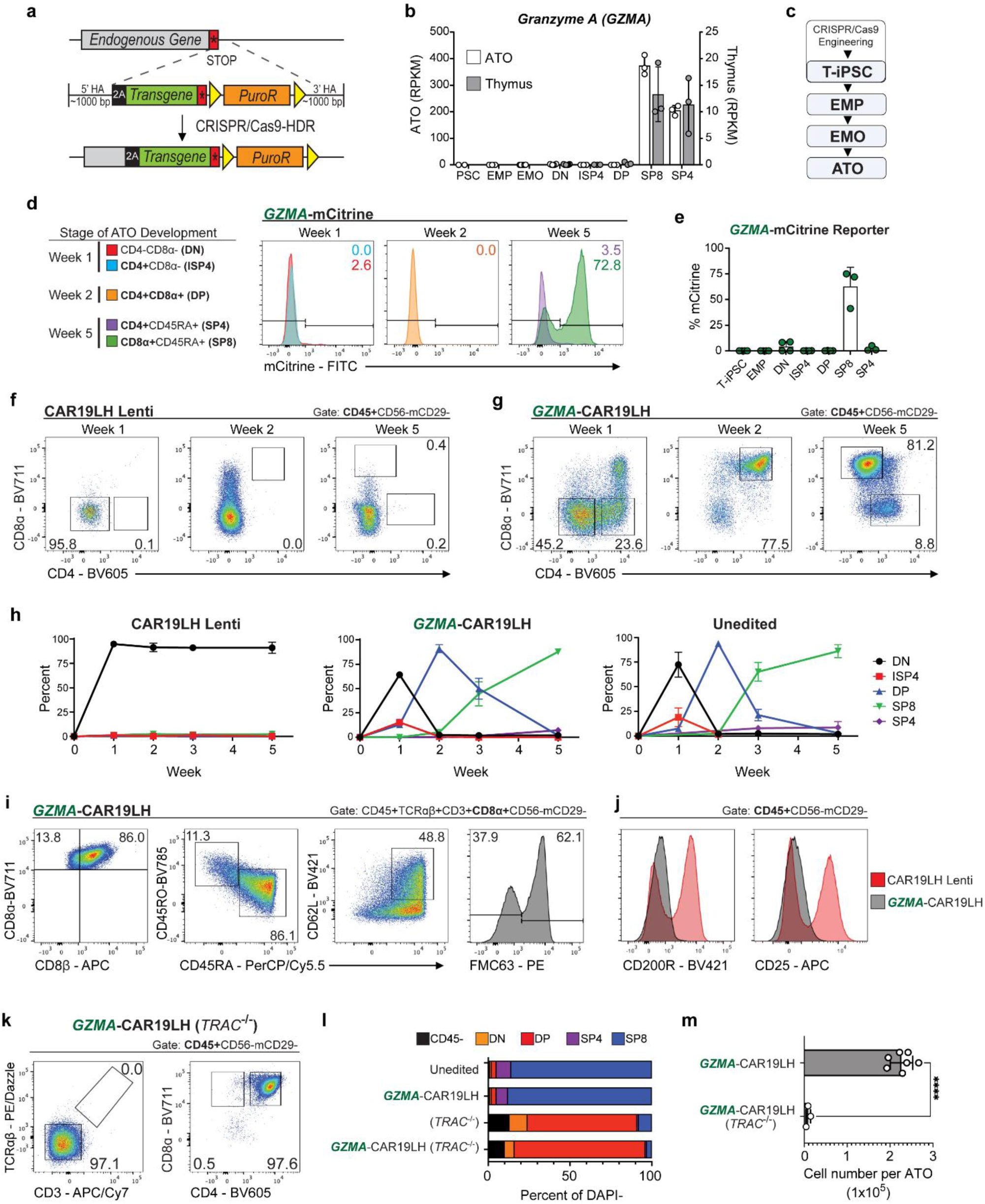
Stage-specific expression of the CAR using the *GZMA* locus rescues conventional T development. **(a)** Schematic of gene editing strategy for the insertion of a transgene into the 3’ UTR of an endogenous gene. CRISPR/Cas9 HDR template was designed to replace the endogenous stop codon (red box) with a 2A ribosome skip sequence, followed by the transgene cDNA in-frame for translation. The template included *loxp*-flanked (yellow triangles) puromycin resistance gene (PuroR) to enrich for edited iPSC clones. 5’ and 3’ homology arms (HA) complementary to the Cas9-mediated cut site were used to direct homologous recombination. **(b)** Bulk RNA-Seq measurements of *GZMA* expression at each stage of T cell development during ATO differentiation (from PSC and Cord Blood) and primary human thymus. Gating strategy for the isolation of each population is as follows: Pluripotent stem cell (PSC: TRA-1-60+SSE.A-4+), embryonic mesodermal progenitors (EMP: CD56+CD326-), embryonic mesodermal organoids (EMO: CD34+CD90+CD43+CD45±), double negative (DN: CD45+CD7+CD1a±), immature single positive CD4+ (ISP4: CD45+CD4+CD8-TCR-CD3-CD7+), double positive (DP: CD4+CD8+CD3-TCRɑβ-), single positive CD8+ (SP8: CD3+CD4-CD8+CD45RA+), and single positive CD4+ (SP4: CD3+CD4+CD8-CD45RA+). Sources for each population are as follows: undifferentiated PSC (left y-axis, white), PSC-derived EMP, EMO, DP, SP8, SP4 (left y-axis, white), CB-ATO derived DN, ISP4 (left y-axis, white), and primary thymus derived DN, ISP4, DP, SP8, SP4 (right y-axis, gray). RPKM = reads per kilo base per million mapped reads. Bulk RNA-seq original data sources derived from GSE116015^19^ and PRJNA741323^20^ (mean ± SD, n = 6 biological replicates for EMO and DN; n = 3 biological replicates for all other cell types). **(c)** Simplified schematic of iPSC differentiation using the Artificial Thymic Organoid (ATO) culture system. CRISPR/Cas9 edited T-iPSCs were differentiated into embryonic mesodermal progenitors (EMP) and then aggregated with MS5-hDLL4 into embryonic mesodermal organoids (EMOs). After 2 weeks of hematopoietic induction, EMOs were re-aggregated with MS5-hDLL4 to generate ATOs. **(d)** Analysis of reporter expression at each T cell stage during differentiation of *GZMA*-mCitrine T-iPSC in ATOs. T cell populations analyzed at weeks 1, 2 and 5 of ATO culture are indicated in the key to the left. mCitrine negative gating was based on unedited T-iPSC ATO control at the same stage of T cell development (data representative of n = 4 independent experiments). **(e)** Summary of reporter data at each stage of *GZMA*-mCitrine T-iPSC differentiation in ATOs (mean ± SD, n = 4 independent experiments). **(f-g)** Analysis of T cell development in ATOs differentiated from **(f)** CAR19LH Lenti and **(g)** *GZMA*-CAR19LH T-iPSCs at indicated time points (data representative of n = 3 independent experiments). **(h)** Kinetics of T cell development during 5 weeks of ATO differentiation using T-iPSC lines shown above each graph. Percent of T cell subsets are of total CD45+CD56-mCD29-DAPI-cells (mean ± SD, n = 3 independent experiments). **(i)** Phenotype and CAR19LH expression (anti-FMC63) in *GZMA*-CAR19LH SP8 T cells analyzed at week 5 of ATO culture (data representative of n = 5 independent experiments). **(j)** Expression of ILC2 markers CD25 and CD200R in CAR19LH Lenti- and *GZMA*-CAR19LH ATO-derived cells analyzed at week 5 (data representative of n = 3 independent experiments). **(k)** Analysis of T cell development in *GZMA*-CAR19LH (*TRAC*^-/-^) ATOs at week 6 of culture (data representative of n = 3 independent experiments). **(l)** Frequencies of T cell subsets at week 6 of ATO culture generated from unedited, *GZMA*-CAR19LH (i.e. *TRAC* intact), *TRAC*-disrupted only control (*TRAC*^-/-^), and *GZMA*-CAR19LH (*TRAC*^-/-^) T-iPSCs, gated on total DAPI-mCD29-cells (mean ± SD, n = 3 independent experiments). **(m)** Number of total cells generated per ATO at week 6 of culture based on manual trypan blue counting (mean ± SD, \*\*\*\**P* < 0.0001 by two-tailed unpaired *t*-test, n = 3 independent experiments for *GZMA*-CAR19LH (*TRAC*^-/-^) T-iPSC; n = 4 independent experiments for *GZMA*-CAR19LH T-iPSC).

Endogenous loci for CAR insertion were identified using bulk RNA sequencing data generated at each stage of T cell differentiation using cells derived from the primary human thymus and ATO cultures^19,20^. As initial proof of concept, the endogenous gene encoding Granzyme A (*GZMA*) was chosen based on low to absent expression throughout differentiation from iPSCs and most critically at the CD4-CD8- (double negative i.e. DN) stage when the fate decision between innate and conventional T lineages occurs; *GZMA* expression was highest in mature, single positive CD8+ (SP8) T cells **(Fig. 1b).**

This editing approach was first validated by the generation of *GZMA* reporter T-iPSCs. The fluorescent reporter mCitrine was inserted at the 3’ UTR of *GZMA* (*GZMA-*mCitrine) and single-cell clones were differentiated using previous published protocols for the ATO culture system^19,21^ **(Fig. 1c and Supplemental Fig. 1a,b).** *GZMA*-mCitrine T-iPSCs underwent normal conventional T cell development, from DN through immature single positive CD4+ (ISP4) at week 1, CD4+CD8+ (double positive i.e. DP) stage by week 2, and underwent positive selection to mature into single positive CD4+ (SP4) and SP8 T cells by week 5 **(Supplemental Fig. 1c-e).** mCitrine reporter expression was absent in T-iPSC prior to differentiation, embryonic mesoderm progenitors (EMP), and T cell precursors; however, onset of reporter expression was observed after the DP stage, particularly in SP8s **(Fig. 1d,e).** mCitrine+ cells were also seen in a small subset of DNs that give rise to innate CD8αα lymphoid cells transiently in early ATO cultures **(Supplemental Fig. 1f).** Notably, the intracellular expression of GZMA protein was confirmed at levels comparable to unedited SP8 T cells **(Supplemental Fig. 1g).** Together, strict stage-specific expression of an exogenous gene during T cell differentiation from iPSCs was achieved by the insertion of a transgene at the 3’ UTR of an endogenous gene.

### *GZMA*-regulated CAR expression allows normal conventional T development to proceed

The same gene editing strategy was next deployed to regulate CAR expression during ATO development. For these studies, a second-generation, anti-CD19 1928z CAR containing a CH2-CH3 spacer in the hinge domain (hereafter CAR19 “long hinge”, i.e. CAR19LH) and known to possess high levels of tonic signaling was tested^7^ **(Supplemental Fig. 2a)**. *GZMA*-CAR19LH T-iPSC clones were generated and differentiated in ATOs in parallel with unedited, parent T-iPSCs (hereafter, “Unedited T-iPSCs”), and T-iPSCs that constitutively expressed CAR19LH via lentiviral transduction (hereafter, “CAR19LH Lenti”).

As previously described^7^, conventional T cell differentiation from CAR19LH Lenti T-iPSCs did not proceed beyond the DN stage **(Fig. 1f).** In contrast, *GZMA*-CAR19LH ATOs showed robust conventional T cell development that mirrored the differentiation of unedited ATOs, with >80% SP8 T cells by week 5 **(Fig. 1g,h).**

*GZMA*-CAR19LH SP8 T cells displayed a conventional (CD8αβ+), mature (CD45RA+) phenotype with 40-50% expressing CD62L **(Fig. 1i).** The frequency of CAR expressing cells mirrored the pattern of *GZMA*-reporter expression (**Fig. 1d,i**) and further increased to >99% upon expansion using CD19+ artificial antigen expressing cells (aAPCs) **(Supplemental Fig. 2b).**

*GZMA*-CAR19LH SP8 T cells lacked expression of ILC2 markers in contrast to cells produced from CAR19LH Lenti T-iPSCs **(Fig. 1j)** and showed antigen-specific production of effector cytokines and CD107a expression **(Supplemental Fig. 2c).**

Together, the delayed, stage-specific expression of a CAR with high tonic signaling overcame diversion towards to ILC2s and resulted in the generation of conventional, functional SP8 CAR T cells from T-iPSCs.

### Disruption of the *TRAC* locus halts T cell differentiation at the DP Stage

To address the risk of alloreactivity, the surface expression of endogenous TCRs was abolished using CRISPR/Cas9 to disrupt exon 1 of the T cell receptor alpha chain constant region (hereafter, *TRAC*^-/-^)^22^ **(Supplemental Fig. 2d,e).** Similar to control *TRAC^-/-^* T-iPSCs, T cell development from *GZMA*-CAR19LH (*TRAC*^-/-^) T-iPSC did not advance beyond the DP stage **(Fig. 1k,l)** and cell yields were significantly reduced compared to *TRAC*-intact *GZMA*-CAR19LH cultures **(Fig. 1m).** As the endogenous *GZMA* gene is not expressed prior to positive selection, *GZMA*-CAR19LH (*TRAC*^-/-^) cells from ATO cultures did not express the CAR transgene **(Supplemental Fig. 2f).** Thus, although *GZMA*-regulated CAR expression prevented the diversion of differentiation to the innate lineages, an alternative approach was required to rescue positive selection in the absence of an endogenous TCR.

### Signaling through CAR at the DP stage rescues positive selection during *TRAC-/-* iPSC differentiation

We next tested if CAR signaling can be used as an alternative mechanism of achieving positive selection. The endogenous *CD8β* locus was chosen to regulate the expression of the CAR based on its absent expression at the immature DN and ISP4 T cell stages but high expression at the DP and SP8 stages **(Fig. 2a).** A *CD8β*-mCitrine reporter T-iPSC line demonstrated the predicted pattern of reporter expression during ATO differentiation **(Fig. 2b).**

**Figure 2:**
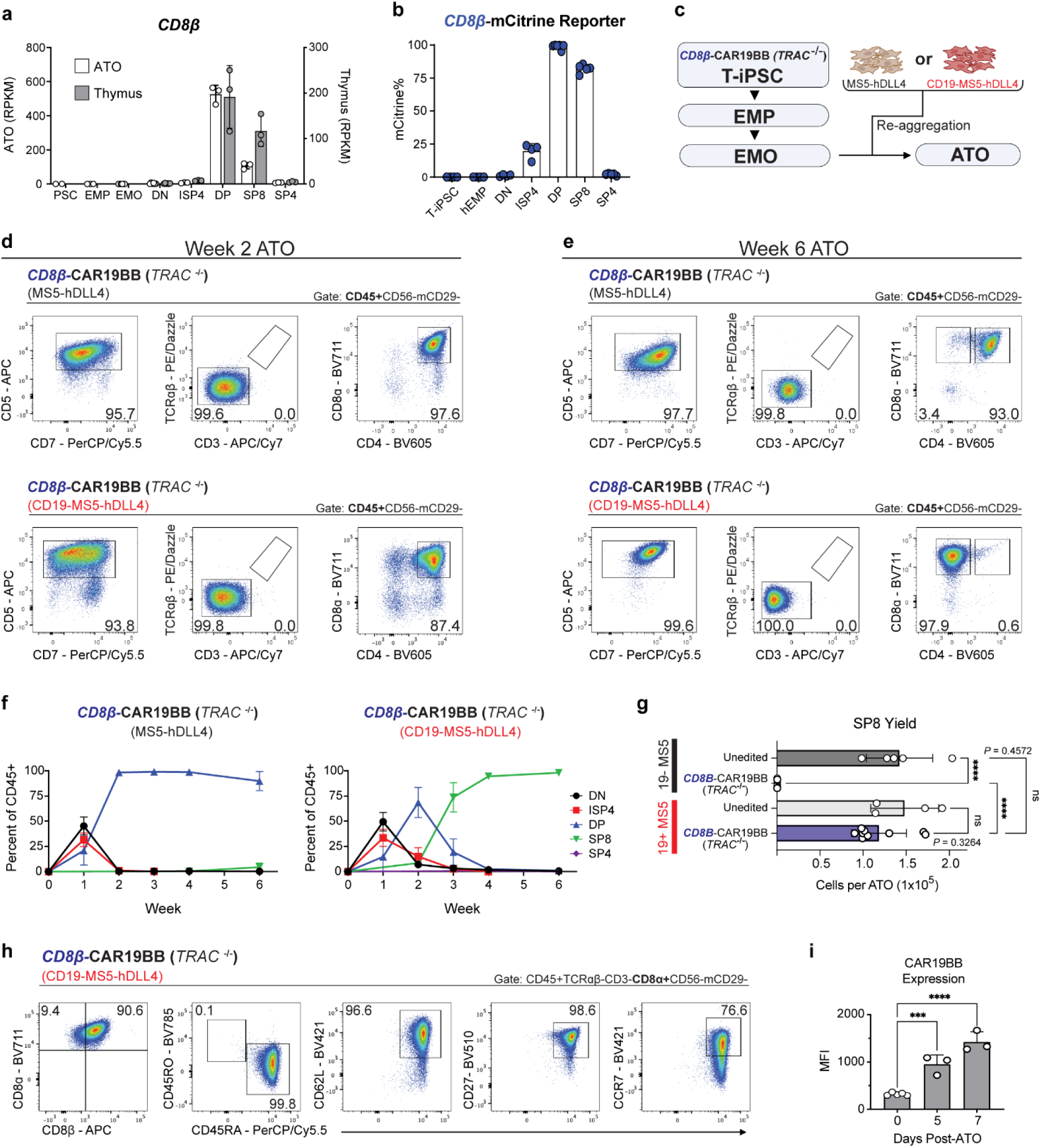
CAR-mediated positive selection using *CD8B-*regulated CAR expression generates CD3-TCRɑβ-SP8 CAR T cells. **(a)** Bulk RNA-Seq measurements of *CD8β* expression at each stage of T cell development during PSC-ATO and CB-ATO differentiation measured as RPKM (left y-axis, white) and primary human thymus (right y-axis, gray). Gating strategy for each population was previously outlined in Figure 1A. RPKM = reads per kilo base per million mapped reads. Bulk RNA-seq original data sources derived from GSE116015^19^ and PRJNA741323^20^ (mean ± SD, n = 6 biological replicates for EMO and DN; n = 3 biological replicates for all other cell types). **(b)** Summary of reporter data at each stage of *CD8β*-mCitrine T-iPSC differentiation in ATOs (mean ± SD, n = 4 independent experiments). **(c)** Simplified schematic of *CD8B-*CAR19BB (*TRAC*^-/-^) T-iPSC differentiation in ATOs. After undergoing hematopoietic induction in EMOs, isolated progenitors were re-aggregated with MS5-hDLL4 or CD19-MS5-hDLL4 to generate ATOs. **(d,e)** Analysis of T cell development in *CD8B-*CAR19BB (*TRAC*^-/-^) ATOs using MS5-hDLL4 (top) or CD19-MS5-hDLL4 (bottom) at **(d)** week 2 and **(e)** week 6 (data representative of n = 6 independent experiments). **(f)** Kinetics of T cell development during 6 weeks of *CD8B-*CAR19BB (*TRAC*^-/-^) T-iPSC differentiation in ATOs generated with MS5-hDLL4 and CD19-MS5-hDLL4. Percent of T cell subsets were of total CD45+CD56-mCD29-DAPI- (mean ± SD, n = 4 independent experiments). **(g)** Mean number of total SP8 T cells generated per ATO at week 6 of culture, based on manual trypan blue counting (mean ± SD, \*\*\*\**P* < 0.0001 by two-tailed unpaired *t*-test, n = 5 independent experiments for unedited T-iPSC, n = 6 independent experiments for *CD8B-*CAR19BB (*TRAC*^-/-^) T-iPSC). **(h)** Phenotype of *CD8B-*CAR19BB (*TRAC*^-/-^) SP8 T cells analyzed at week 6 of CD19-expressing ATO cultures (data representative of n = 6 independent experiments). **(i)** CAR19BB expression (Mean Fluorescence Intensity (MFI) of anti-FMC63) in *CD8B-*CAR19BB (*TRAC*^-/-^) SP8 T cells analyzed at week 6 of CD19-expressing ATO cultures (Day 0), and 5 and 7 days after isolation from ATOs (post-ATO) (mean ± SD, \*\*\**P* = 0.0003, \*\*\*\**P* < 0.0001 by two-tailed unpaired *t*-test, n = 5 independent experiments for post-ATO day 0 and n = 3 independent experiments for days 3 and 5).

A second generation anti-CD19 4-1BB (19BBz) CAR^7^ was inserted into the *CD8β* locus of *TRAC*^-/-^ T-iPSCs [hereafter *CD8B-*CAR19BB (*TRAC*^-/-^)] **(Supplementary Fig. 3a).** To provide the CAR cognate antigen during development, CD19 was constitutively expressed in MS5-hDLL4 via lentiviral transduction (CD19-MS5-hDLL4) **(Supplementary Fig. 3b)**. *CD8B-*CAR19BB (*TRAC*^-/-^) T-iPSCs were differentiated into EMOs using the standard MS5-hDLL4 stroma, but CD19-MS5-hDLL4 was substituted for the generation of ATOs; ATOs generated with parent MS5-hDLL4 (without expression of CD19 antigen) served as controls **(Fig. 2c).**

In both the presence and absence of the CD19 antigen, *CD8B-*CAR19BB (*TRAC*^-/-^) ATO cultures reached the DP T cell precursor stage by week 2 without diversion to ILCs **(Fig. 2d and Supplemental Fig. 3c).** By week 6, *CD8B-*CAR19BB (*TRAC*^-/-^) ATOs generated in the absence of CD19 antigen remained at the DP stage with little, if any, positive selection **(Fig. 2e,f)**. In contrast, robust positive selection was achieved in CD19-expressing ATOs, as CD3-TCRɑβ-SP8 T cells became the dominant population in the culture **(Fig. 2e,f).** Total SP8 yield per ATO from *CD8B-*CAR19BB (*TRAC*^-/-^) T-iPSCs was comparable to ATOs generated using unedited T-iPSCs **(Fig. 2g)**.

Closer analysis of *CD8B-*CAR19BB (*TRAC*^-/-^) SP8 T cells revealed a conventional CD8αβ phenotype with a highly uniform “naïve” (Tn-like) phenotype; over 90-96% of SP8 cells expressed CD45RA, CD62L and CD27, and over 75% expressed high levels of CCR7 **(Fig. 2h and Supplemental Fig. 3d).** Cell surface CAR expression was detectable at low levels in CD19-expressing ATOs but increased once cells were isolated from ATOs and rested in the absence of CD19 antigen **(Fig. 2i and Supplementary Fig. 3e).**

Importantly, CAR-mediated positive selection and the production of naïve CAR T cells was not limited to a particular CAR design. TRAC^-/-^ T-iPSCs similarly modified to express the high tonic signaling CAR19LH in DPs [*CD8β*-CAR19LH (*TRAC*^-/-^)] achieved comparable levels of positive selection in the presence of CD19, with uniform generation of Tn-like SP8 cells **(Supplemental Fig. 4a-d).** CAR-mediated positive selection with production of CD45RA+CD62L+CCR7+ SP8 T cells was also observed by targeting CD20 and using an anti-CD20 CAR **(Supplemental Fig. 4e).** In addition, removal of MHC class 1 expression through the deletion of *B2M* did not impair Tn-like SP8 development from *CD8B-*CAR19BB (*TRAC*^-/-^) T-iPSCs **(Supplemental Fig. 4f).**

### Naïve-like T cells differentiate into effector memory T cells during post-ATO stimulation

During normal conventional T cell development, cells exit the thymus as Tn cells and can further differentiate to effector memory T cells (Tem) upon encountering antigens in the periphery. To test if the Tn-like CAR T cells generated from ATO cultures can generate Tem cells, CD45RA+CD62L+CCR7+ SP8 T cells were first isolated from *CD8β*-CAR19BB (*TRAC*^-/-^) week 5 ATOs **(Supplemental Fig. 5a)** and stimulated using CD19+ aAPCs. Over the course of a 7-day post-ATO stimulation, CD45RO expression in SP8 cells progressively increased with a reciprocal decrease in CD45RA expression; CD62L and CCR7 were downregulated within the first 3 days, while CD27 expression was maintained **(Fig. 3a)**. The CD45RO+CD62L-CCR7-CD27+ Tem-like phenotype remained largely unchanged during the second and subsequent stimulations **(Supplemental Fig. 5b,c).** Importantly, expression of classical exhaustion markers on *CD8β*-CAR19BB (*TRAC*^-/-^) SP8 T cells was low to absent during development in the ATOs and did not significantly increase after undergoing multiple cycles of post-ATO stimulations **(Supplemental Fig. 5d,e).**

**Figure 3:**
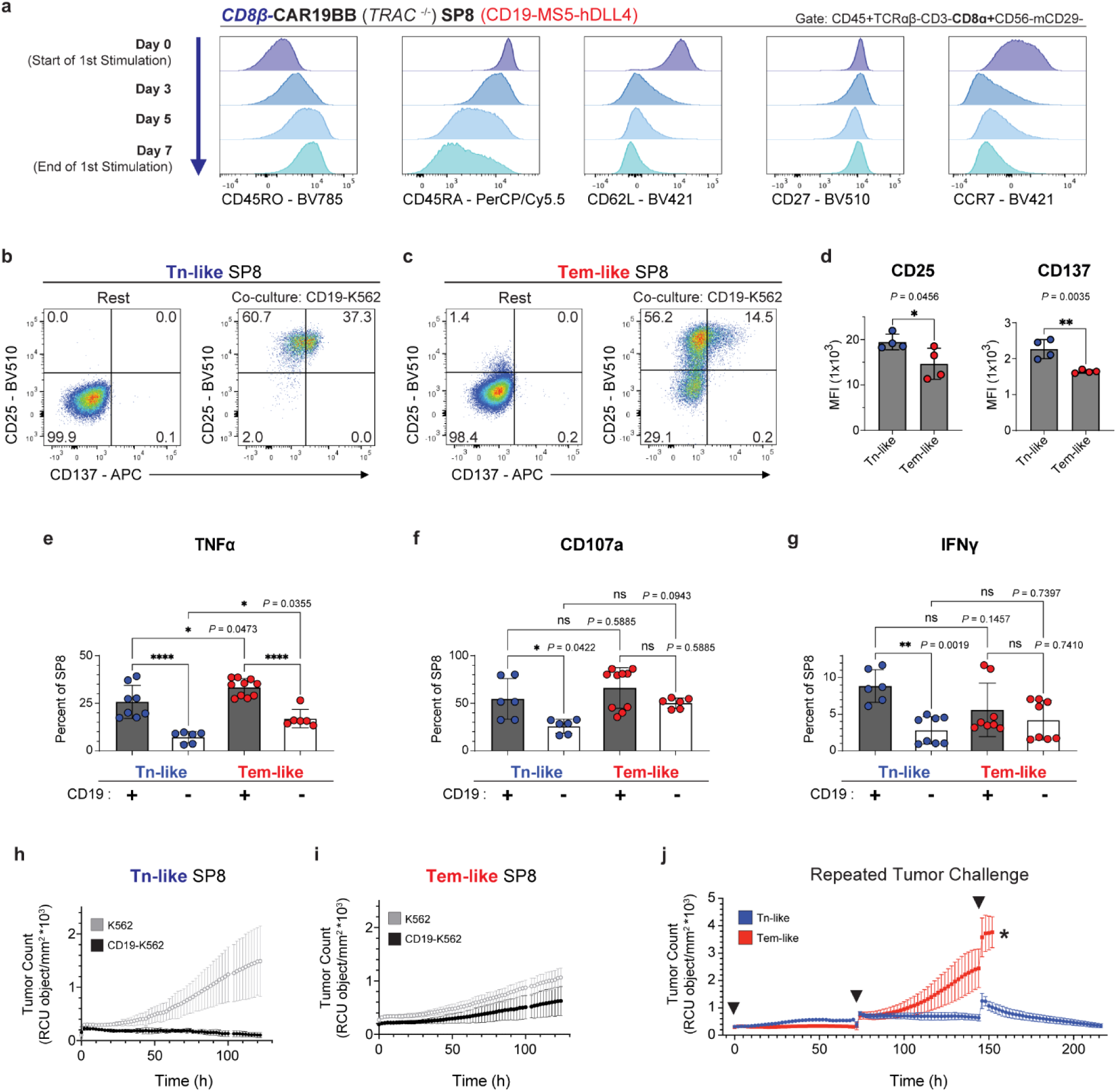
Impact of T cell stimulation on SP8 CAR T cell phenotype and function. **(a)** Change in SP8 T cell phenotype during aAPC-mediated stimulation. Isolated *CD8B-*CAR19BB (*TRAC*^-/-^) SP8 T cells (at week 5 of CD19-expressing ATO cultures) were co-cultured with CD19-K562-CD137L aAPCs and phenotype was evaluated at 3, 5, and 7 days of aAPC stimulation (data representative of n = 2 independent experiments). **(b,c)** CD25 and CD137 (4-1BB) expression in **(b)** Tn-like and **(c)** Tem-like SP8 T cells in response to 24h co-culture with CD19-K562 compared to baseline (Rest). All SP8 T cells are gated CD45+CD7+CD8ɑ+CD56-mCD29-DAPI- (data representative of n = 4 independent experiments). **(d)** CD25 and CD137 (4-1BB) expression (MFI) in Tn-like and Tem-like SP8 T cells in response to 24h co-culture with CD19-K562 (mean ± SD, \**P* < 0.05 by two-tailed unpaired *t*-test, n = 4 independent experiments). **(e-g)** Quantification of **(e)** TNFα, **(f)** CD107a, and **(g)** IFNγ expressing Tn-like (blue circles) and Tem-like (red circles) SP8 T cells in response to 6h co-culture with CD19-K562 (black bar) and antigen negative K562 controls (white bar) (mean ± SD, \*\*\*\**P* < 0.0001 by ordinary one-way ANOVA followed by Tukey’s multiple comparison test, n = 4 independent experiments). **(h,i)** Incucyte tumor challenge assay measuring the growth of K562 (gray) or CD19-K562 (black) tumor cells co-cultured with **(h)** Tn-like or **(i)** Tem-like SP8 T cells at a Target:T cell ratio of 1:1 (data representative of n = 2 independent experiments). **(j)** Repeated tumor challenge measuring the growth of CD19-K562 tumor cells (by Incucyte assay) co-cultured with Tn-like or Tem-like SP8 T cells at a Target:T cell ratio of 1:1. 15,000 live tumor cells were added to the cultures at time shown (black triangles). Black asterisk denotes culture failure due to tumor overgrowth (data representative of n = 2 independent experiments).

### CAR T cell function is impaired upon loss of naïve-like phenotype

To determine if there was any functional significance of the change in T cell phenotype during culture, functional assays of *CD8β*-CAR19BB (*TRAC*^-/-^) SP8 T cells were performed before stimulation (when >80% of cells were Tn-like) and after undergoing two, 7-day stimulations using aAPCs (when >90% were Tem-like) **(Supplemental Fig. 6a).**

Co-culture with each T cell population with CD19+ tumor cells for 24 hours showed significantly higher activation in Tn-like than Tem-like SP8s (based on CD25 and CD137 upregulation) **(Fig. 3b-d**). Although both Tn- and Tem-like T cells showed antigen-specific TNFα production (**Fig. 3e**), only Tn-cells demonstrated antigen-specific CD107a and IFNγ upregulation **(Fig. 3f,g and Supplemental Fig. 6b-e)**; however, the responses seen in Tem-like cells were antigen-independent **(Fig. 3f,g and Supplemental Fig. 6b-e)**.

The difference in antigen-specificity between the two phenotypes became more evident using an *in vitro* cytotoxicity assay; antigen-specific clearance of CD19+ tumor cells was restricted to Tn-like SP8 T cells, while Tem-like SP8 T cells showed cytotoxicity, regardless of the presence of the CD19 antigen **(Fig. 3h,I and Supplemental Fig. 6f,g).** Under repeated challenge, Tn-like cells demonstrated complete clearance of tumor cells, whereas Tem-like SP8s lost the ability to control tumor after the first challenge **(Fig. 3j and Supplemental Fig. 6h).** Taken together, these experiments suggest that CAR-expressing Tn-like SP8 cells generated in ATOs produced more potent and antigen-specific responses compared to cells that have undergone differentiation to Tem-like SP8s.

### Development and maintenance of the naïve phenotype is dependent on generation in the 3-D ATO context

The production of SP8 CAR T cells from iPSCs has been accomplished by others by isolating DP cells that have been generated in monolayer or stroma-free systems and then stimulating those isolated DP T cells with aAPCs or anti-CD3/CD28 beads^9,12,23^. Although SP8 T cells produced with these approaches were shown to be CD8αβ+, the uniform Tn-like phenotype observed in *CD8β*-CAR19BB (*TRAC*^-/-^) SP8 T cells has not to our knowledge been previously reported.

To explore the influence of the ATO microenvironment on CAR-mediated positive selection, SP8 T cells produced from *CD8β*-CAR19BB (*TRAC*^-/-^) ATO cultures (in which CAR stimulation was provided by the CD19-expressing MS5-hDLL4 stroma) were compared to SP8 T cells produced by antigen stimulation of ATO-derived DP precursors (isolated at week 2 of culture) using aAPCs. The output and phenotype of T cells from each condition were evaluated at the same relative period of differentiation and stimulation (**Fig 4a**).

**Figure 4:**
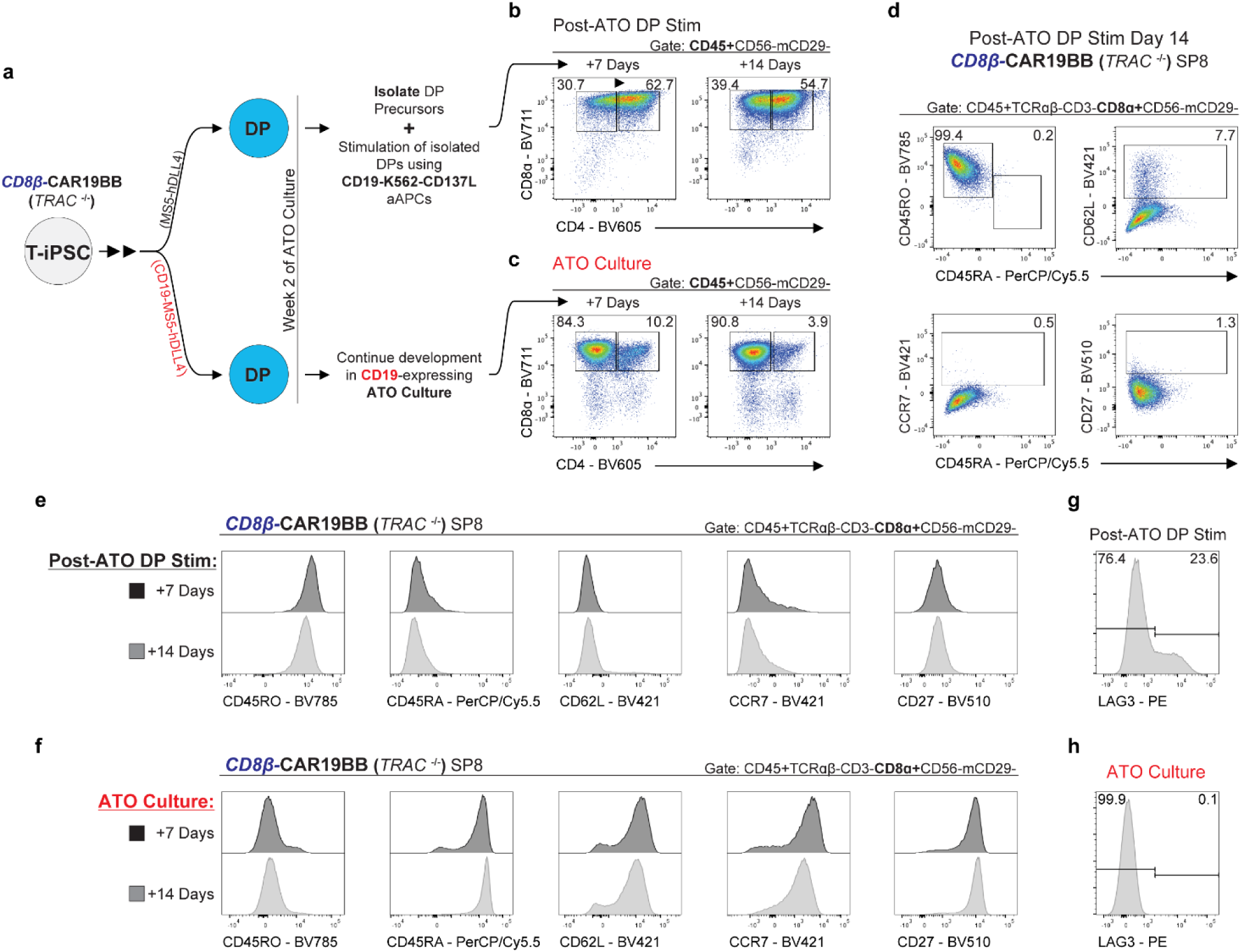
Induction and maintenance of the naïve phenotype is dependent on CAR signaling in the ATO context. **(a)** Simplified schema comparing two methods to induce CAR-mediated positive selection. *CD8B-*CAR19BB (*TRAC*^-/-^) T-iPSCs underwent ATO differentiation to generate DP precursors in the presence or absence of CD19 antigen. At week 2 of ATO culture, DP precursors were isolated from CD19-negative ATOs and co-cultured using CD19-K562-CD137L aAPCs for 14 days. In parallel, CD19-expressing ATOs remained unperturbed in ATO culture for a further 14 days. Both conditions were analysed at day 7 and day 14 of culture post-DP development. **(b,c)** Analysis of SP8 T cell development after **(b)** stimulation of isolated ATO-derived DP precursors using CD19+ aAPCs and **(c)** continued CD19-expressing ATO cultures, evaluated at the same relative time points indicated above (data representative of n = 2 independent experiments). **(d)** Phenotype of *CD8B-*CAR19BB (*TRAC*^-/-^) SP8 T cells generated after 14-day stimulation of isolated ATO-derived DP precursors using CD19+ aAPCs (data representative of n = 2 independent experiments). **(e,f)** Phenotype of *CD8B-*CAR19BB (*TRAC*^-/-^) SP8 T cells after **(e)** stimulation of isolated ATO-derived DP precursors using CD19+ aAPCs and **(f)** continued CD19-expressing ATO cultures at day 7 (black) and day 14 (grey) of culture post-DP development (data representative of n = 2 independent experiments). **(g,h)** LAG3 expression in *CD8B-*CAR19BB (*TRAC*^-/-^) SP8 T cells generated after **(g)** stimulation of isolated ATO-derived DP precursors using CD19+ aAPCs and **(h)** continued CD19-expressing ATO cultures at day 14 of culture. SP8 T cells are gated CD45+TCRɑβ-CD3-CD8ɑ+CD56-mCD29-DAPI- (data representative of n = 2 independent experiments).

14 days after DP isolation, aAPC-stimulated cultures showed incomplete positive selection, with only 40% TCRɑβ-CD3-CD8αβ+ T cells and persistence of ∼50% DPs **(Fig. 4b)**. By contrast, 14 days of continued ATO culture produced >90% SP8 T cells **(Fig. 4c)**. Notably, SP8 CAR T cells generated from aAPC-mediated stimulation of isolated DP T cells showed a predominant Tem-like phenotype with high expression of CD45RO but little to no expression of CCR7, CD62L or CD27 (**Fig. 4d,e**), similar to the phenotype of post-ATO stimulated SP8s **(Fig. 3a).** Conversely, SP8 CAR T cells derived from CAR signaling during continuous ATO culture showed uniform expression of CD45RA, CD62L, CCR7, and CD27 at the onset of SP8 development (**Fig. 4f**), which remained high and unchanged over an additional 4 weeks of ATO culture **(Supplemental Fig. 7a)**. In addition, 14 days of isolated DP stimulation by aAPCs resulted in the upregulation of the exhaustion marker LAG3, while expression was absent in continued ATO cultures **(Fig. 4g,h).** These experiments suggest unique interactions with the engineered CD19-MS5-hDLL4 stroma, in concert with the 3-D organoid microenvironment, are required to induce and preserve the Tn-like phenotype.

### Transcriptional comparisons between TCR- and CAR-mediated T cell development

To explore the potential mechanisms mediating the Tn-like phenotype seen with CAR-mediated positive selection, single-cell RNA and ATAC sequencing was performed on unsorted cells from CD19-expressing ATO cultures generated from *CD8B-*CAR19BB (*TRAC*^-/-^) T-iPSCs and ATO cultures generated from unedited T-iPSCs (in which positive selection is mediated through TCR-pMHC self-selection). Cultures were harvested at week 3, allowing the capture of both DP precursors and SP8 T cells **(Supplemental Fig. 7b,c).** A multimodal weighted nearest neighbor (WNN) uniform manifold approximation and projection (WNN_UMAP) visualization that integrated both single-cell RNA and ATAC libraries confirmed the robust generation of SP8 T cells and DP precursors from both edited *CD8B*-CAR19BB (TRAC^-/-^) and unedited T-iPSCs **(Fig. 5a and Supplemental Fig. 8a-c).**

**Figure 5:**
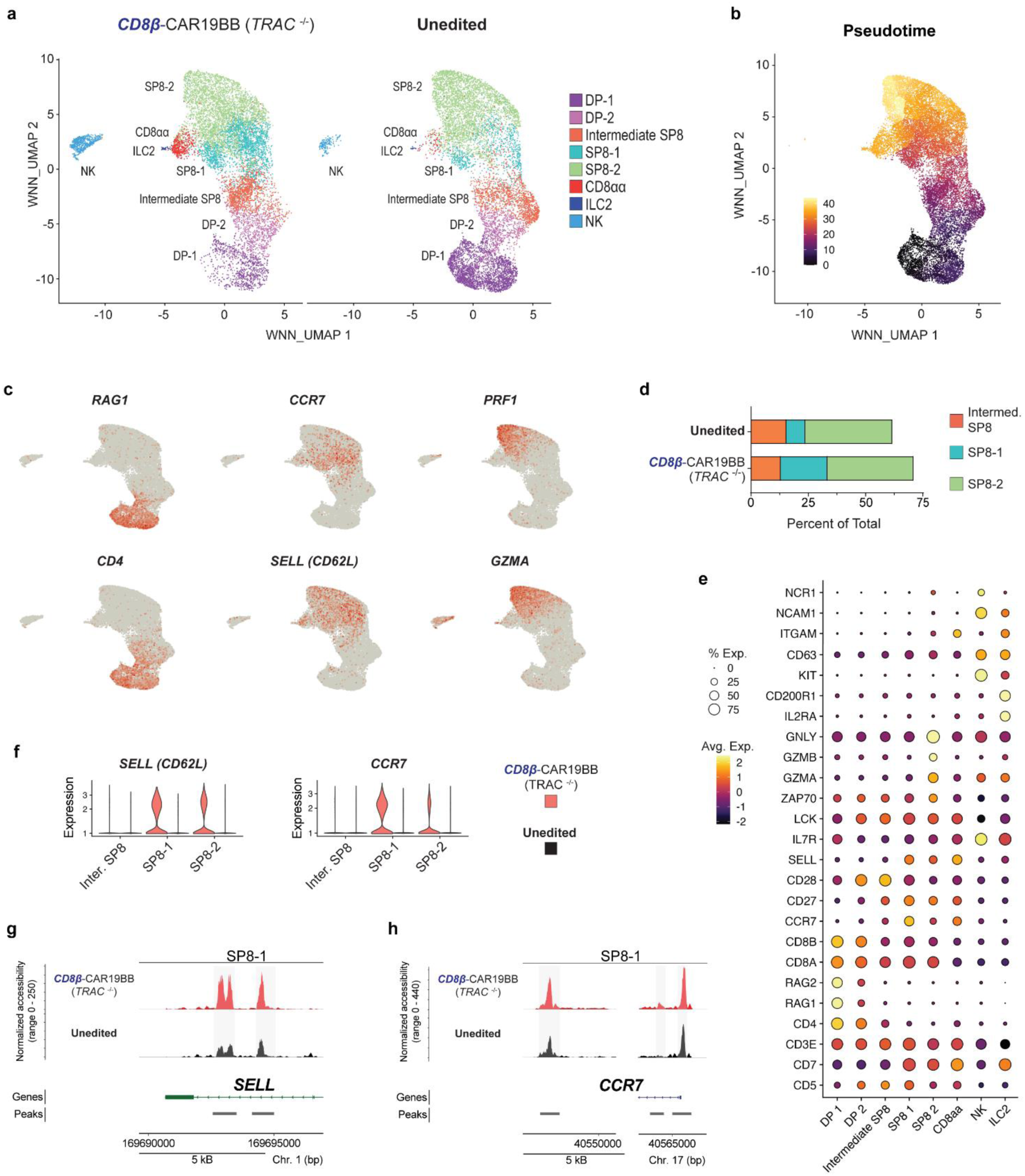
Multimodal analysis of endogenous TCR- and CAR-mediated development in ATO cultures. **(a)** WNN UMAP visualization of annotated populations from unedited (right) and *CD8B-*CAR19BB (*TRAC*^-/-^) (left) ATOs sequenced at week 3 of culture. UMAP was generated using the integration of both RNA and ATAC libraries. **(b)** Combined multimodal UMAP visualizing pseudotime trajectory represented as a heatmap. DP-1 cluster was chosen as the root of pseudotime trajectory. NK cells were excluded from the analysis. **(c)** Expression of selected genes mapped over WNN UMAP visualization. **(d)** Percent of individual SP8 clusters within total cells sequenced from unedited and *CD8B-*CAR19BB (*TRAC*^-/-^) ATOs. **(e)** Average expression of lineage defining genes for clusters identified in cells sequenced from both unedited and *CD8B-*CAR19BB (*TRAC*^-/-^) ATOs. Circle size represents the percentage of cells expressing genes and heatmap represents average expression. **(f)** Violin plots of *SELL* and *CCR7* gene expression in SP8 clusters identified from unedited (black) and *CD8B-*CAR19BB (*TRAC*^-/-^) (red) ATOs. **(g,h)** Genome browser tracks of **(g)** *SELL* and **(h)** *CCR7* chromatin accessibility comparing both samples within the SP8-1 cluster. ATAC-seq peaks are normalized between the samples.

Further sub-clustering of T cell subsets resulted in two DP sub-clusters (DP-1 and -2), an “intermediate” SP8 cluster, and two SP8 sub-clusters (SP8-1 and SP8-2) (**Fig 5a**). These cluster identities were confirmed using pseudotime analysis, which showed linear trajectory of T cell development originating from the DP-1 and ending at the SP8-2 cluster **(Fig. 5b).** Mature T cells (SP8-1 and SP8-2) and T precursors (DP-1, DP-2, and intermediate SP8) accounted for 88% and 97% of identified cells in *CD8B-*CAR19BB (*TRAC*^-/-^) and unedited ATOs, respectively **(Supplementary Fig. 8d).**

Projection of canonical T cell gene expression mapped on the multimodal UMAP showed high expression of *RAG1* and *CD4* in DP-1 and DP-2 clusters, while intermediate SP8 showed lower levels of *CD4* and no expression of *RAG1* or mature T cell markers, suggestive of the latter subset having recently undergone positive selection **(Fig. 5c and Supplemental Fig. 8e).** Within the mature SP8-1 and SP8-2 clusters, the expression of naïve makers (i.e. *CCR7, SELL)* was primarily restricted to SP8-1, while effector gene expression (i.e. *PRF1, GMZA*) was exclusively restricted to SP8-2 **(Fig. 5c).** Differential gene analysis^24^ revealed SP8-1 represented a resting, naïve SP8 population (**Supplemental Fig. 8f**) that was higher in both the number and proportion of total cells produced using CAR-mediated positive selection, compared to unedited T-iPSCs **(Fig. 5d and Supplemental Fig. 8d).** By contrast, SP8-2 cells represented an activated, effector SP8 population with high expression of genes associated with immune response activation and cytokine production **(Supplemental Fig. 8g).** A small frequency of non-T populations was also generated from both lines, including NK (*NCAM1, KIT, NCR1)*, ILC2 (*CD200R1, IL2RA)*, and CD8ɑɑ innate cells **(Fig. 5e).**

Interestingly, differences in gene expression and chromatin regulation were found within the SP8 clusters based on the T-iPSC line from which they were generated. Both SP8-1 and SP8-2 clusters from *CD8B*-CAR19BB (TRAC^-/-^) ATOs expressed the naïve genes *SELL* and *CCR7*, while expression was low to absent in all unedited SP8 T cells (**Fig. 5f).** Consistent with the gene expression, *CD8B-*CAR19BB (*TRAC*^-/-^) T cells showed greater chromatin accessibility at both naïve *SELL* and *CCR7* loci compared to unedited T cells (**Fig. 5g,h)**.

### CAR-mediated positive selection drives key factors that promote naïve development

The data above demonstrated TCR- and CAR-mediated positive selection result in differences in the frequency and transcriptional profile of T cells within the mature SP8 sub-clusters. To identify the potential mediators that drive these changes, we shifted our analysis further upstream to the precursor stages at which positive selection signals originate (i.e. DP-1 through intermediate SP8).

Analysis of differentially expressed genes within each precursor cluster revealed a common group of transcription factors that were upregulated in *CD8B-*CAR19BB (*TRAC*^-/-^) T cells when compared to unedited T cells at each stage. The transcription factor *BACH2,* which suppresses the differentiation of naïve T cells^25,26^, was significantly upregulated in DP-1, DP-2, and intermediate SP8 and persisted in SP8-1 T cells generated from *CD8B-*CAR19BB (*TRAC*^-/-^) iPSCs **(Fig. 6a)**.

**Figure 6:**
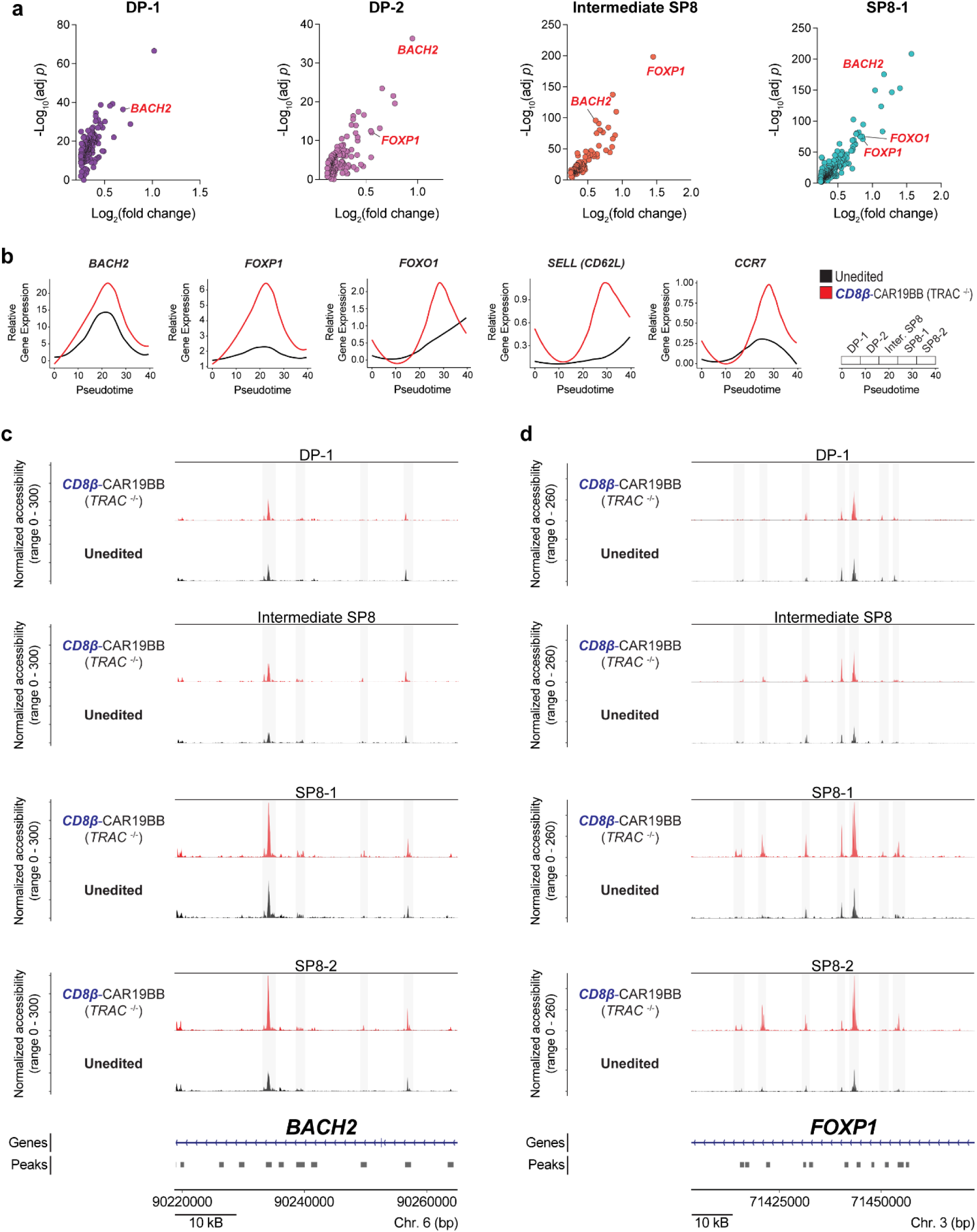
Analysis of T cells and precursors during TCR- or CAR-mediated positive selection. **(a)** Comparison of gene expression within the defined cluster between *CD8B-*CAR19BB (*TRAC*^-/-^) and unedited T cells. All genes with positive log_2_(fold change) > 0.25 in *CD8B-*CAR19BB (*TRAC*^-/-^) T cells compared to unedited T cells are shown **(Extended data 1)**. **(b)** Relative gene expression of select genes across pseudotime in unedited (black) and *CD8B-*CAR19BB (*TRAC*^-/-^) (red) T cells. **(c,d)** Genome browser tracks of **(c)** *BACH2* and **(d)** *FOXP1* chromatin accessibility comparing *CD8B-*CAR19BB (*TRAC*^-/-^) (red) and unedited (black)T cells within DP-1, intermediate SP8, SP8-1 and SP8-2 clusters. ATAC-seq peaks are normalized between the samples.

Similarly, the transcription factor *FOXP1,* which maintains quiescence of naive T cells^27,28^, was significantly upregulated from the DP-2 stage throughout differentiation into SP8-1 T cells **(Fig. 6a)**. Pseudotime analysis also demonstrated the higher relative gene expression of *BACH2* and *FOXP1*, as well as *FOXO1*—another transcription factor that directly regulates naïve T cell development^29–31^—in *CD8B-*CAR19BB (*TRAC*^-/-^) T cells throughout differentiation, compared to unedited T cells **(Fig. 6b).** Notably, the peak expression of these transcription factors immediately proceeded or correlated directly with the expression of naïve genes *SELL* and *CCR7* **(Fig. 6b).**

Chromatin accessibility of the same, key transcription factors further reinforced the differences between *CD8B-*CAR19BB (*TRAC*^-/-^) and unedited cells within each individual T cell cluster. Precursor T cells from *CD8B-*CAR19BB (*TRAC*^-/-^) ATO cultures showed relatively similar accessibility at the *BACH2* locus compared to unedited T cells (**Fig. 6c**). However, upon reaching the SP8 stages, accessibility of the *BACH2* locus increased and remained two- and three-fold more open than unedited cells in both SP8-1 and SP8-2 clusters **(Fig. 6c)**. Higher chromatin accessibility was also seen at the *FOXP1* locus in *CD8B-*CAR19BB (*TRAC*^-/-^) T cells in both SP8 clusters **(Fig. 6d)**.

## Discussion

The unique potential of iPSCs for genetic engineering and limitless expansion of qualified single-cell clones provide the underlying rationale that spurs exploration of stem cell technology as an alternative cellular source for immunotherapy. However, the impact of gene modifications in iPSCs on subsequent T cell differentiation requires new considerations of manufacturing design that do not exist for autologous CAR T cells.

Earlier attempts to generate conventional CAR T cells from iPSCs have identified three key considerations: timing of CAR expression during differentiation^6,7^, level of CAR expression^7,11^, and degree of CAR tonic signaling^7,9,12^. We and others have shown that constitutive expression of a high tonic-signaling CAR skews differentiation towards the innate lineage or blocks T cell development altogether^6,7,9^. Though the degree of innate lineage diversion can be reduced by simply decreasing CAR expression, a low level of CAR expression that permits conventional T cell development may not be sufficient to induce optimal antigen-specific T cell function and response^7,32^. One approach to overcome such problems has been to introduce the CAR transgene after *in vitro* T cell differentiation, producing so-called “iCAR T cells”^2,9^. However, this method loses the inherent logistical advantages of introducing all gene modifications at the self-renewing iPSC stage.

More recently, in an attempt to prevent both early CAR and endogenous TCR expression, van der Stegen et. al inserted the CAR transgene into the *TRAC* locus^12^. However, attenuation of CAR tonic signaling by selectively mutating CD3ξ immunoreceptor tyrosine activation motifs (ITAMs) was still required to avoid lineage diversion^12,33^. Alternatively, we and others have also shown that alternative co-stimulatory domains to modulate signaling can prevent CAR-mediated diversion of some CARs^7,9^. However, modifications of the CAR design can unpredictably impact CAR T cell function and persistence^34,35^, and ultimately impact T cell differentiation, thereby limiting the number of CAR designs that might be useful for iPSC-derived therapies.

By contrast, we show that *CD8β-*mediated regulation of CAR expression, combined with the introduction of the cognate antigen during development, bypassed early innate lineage diversion, while simultaneously inducing highly efficient positive selection in the absence of endogenous TCR expression. Importantly, similar levels of CAR-mediated positive selection were achieved irrespective of the degree of tonic signaling, costimulatory factor or choice of target antigen.

To our knowledge, our work is the first to generate T-iPSC-derived, TCRαβ-CD3- CD8αβ+ CAR T cells with high and uniform expression of the key cell surface markers that define a naïve state. We also highlight the functional consequences associated with the naïve phenotype, including significantly improved antigen-specific activation, cytokine production, and improved tumor control *in vitro*. Our findings are consistent with studies that demonstrate CAR T products derived from peripheral blood Tn or stem cell memory T cells (Tscm) provide a greater and more durable anti-tumor response compared to more terminally differentiated CAR T cells^36,37^. Of note, direct functional comparisons between naïve and effector memory T cells using iPSC-derived T cells have not been previously reported.

Interestingly, the induction and maintenance of the naïve phenotype from T-iPSCs was highly dependent on the context by which developing T cells achieved positive selection. We show here that antigen stimulation of isolated DP precursors (an approach commonly used to overcome the inefficient positive selection seen in monolayer and stroma-free systems^9,12,23^) did not generate the Tn-like T cells produced in ATOs. This context-dependent induction of a naïve phenotype suggests other essential interactions unique to the ATO microenvironment are necessary, which may include endogenous signals from the accessory stromal cells or the intercellular interactions in 3-D culture^38,39^.

Single-cell multi-omics revealed that—compared to endogenous TCR-mediated positive selection—stage-specific CAR-mediated positive selection induced higher expression and chromatin accessibility of the transcription factors *BACH2, FOXP1, and FOXO1*. Notably, each transcription factor plays a critical role during the DP to SP transition and the maintenance of naïve T cells. Specifically, both *BACH2* and *FOXP1* suppress the expression of effector memory-related genes and promote naïve T cell quiescence^25–28^. Similarly, *FOXO1* promotes naïve T cell survival by regulating IL-7Rɑ expression and indirectly controls the expression of *SELL* and *CCR7*^29,30^. Both increased gene expression and higher chromatin accessibility of these genes were first observed at the precursor stages where positive selection signals occur and became more pronounced as cells transitioned to SP8s. The persistent and combined activity of these transcription factors may explain the higher expression and chromatin accessibility of *SELL* and *CCR7* from the onset of positive selection to the maturation of *CD8β*-CAR19BB (*TRAC*^-/-^) SP8 T cells. The differential regulation of naïve genes may also be explained by the unique protein-protein interactions and signalosomes, including TRAF-mediated complexes and TAK1, that were recently found to be recruited only during CAR-mediated signaling (and not TCR signaling)^40^. Future work will continue to identify the upstream signaling networks that directly link CAR signaling to the activity of these transcription factors and ultimately to the efficient induction of the naïve T program.

In summary, we present a method to efficiently generate iPSC-derived, TCR-less CAR T cells through stage-specific signaling and positive selection. This method can likely be applied to any type of CAR inserted into self-renewing, undifferentiated iPSCs that have also been edited to remove endogenous TCR expression. Beyond the regulation of CAR expression, the stage-specific expression system could be modified to express different classes of biologically active transgenes (i.e. cytokines, transcription factors) to further enhance T cell function and response. Deploying a developmentally sensitive approach will be critical to realizing the full potential for “off-the-shelf” stem cell derived cellular therapies.

## Materials and Methods

### Cell lines

The MS5-hDLL4 and CD19-MS5-hDLL4 cell lines were generated in our laboratory as previously described^7,21^. CD20-MS5-hDLL4 cell line was generated by transduction of MS5-hDLL4 with a lentiviral vector encoding for CD20 and sorting via FACS using anti-CD20 antibody. aAPCs used for T cell expansion and *in vitro* assays were generated by the transduction of K562-CD19 with a lentiviral vector encoding for CD137L and mKate2-NLS respectively. Transduced cells were purified by FACS using an anti-CD137 antibody or sorting for mKate2+.

### Human induced pluripotent stem cells lines

The UCLA human Embryonic Stem Cell Research Oversight Committee approved all protocols for the use of T-iPSCs for this study. T-iPSC lines (Cedars-Sinai iPSC Core, Los Angeles, CA) were maintained using Matrigel-coated 6-well plates (Growth Factor Reduced Matrigel Matrix; Cat: 356231, Corning) in mTeSR™ Plus complete medium (Cat: 100-0276, Stem Cell Technologies). Culture medium was changed daily. Upon reaching ∼80% confluency, cultures were dissociated into single-cell suspension using TrypLE™ Express (Cat: 12604-013, Thermo Fisher Scientific) and plated at a minimum density of 1.5 x 10^5^ cells/well of a Matrigel-coated 6-well plate in mTeSR™ with Y-27632 dihydrochloride (10 µM, cat: 1254, Tocris Bioscience), which was removed from culture medium after 1 day. Single-cell clones isolated after CRISPR/Cas9-editing were evaluated for chromosomal abnormalities, including G-band karyotyping (Cedars-Sinai Biomanufacturing Center) and genomic instability analysis kit (Cat: 07550, Stem Cell Technologies).

### Lentiviral vector packaging

The full-length coding sequences of *CD20, CD137L,* and *mKate2-NLS* were synthesized as gene blocks (Integrated DNA Technologies) and cloned into third-generation lentiviral vector backbone pCCL-c-MNDU3 (gift from Donald Kohn, University of California, Los Angeles). Anti-CD19 CAR19LH (scFv: FMC63) was designed and generated as previously described^7^. CAR19LH (1928z) features FMC63 scFV, IgG4 hinge, CH2-CH3 spacer, CD28 transmembrane domain, CD28 co-stimulatory domain, and CD3ζ intracellular signaling domain^7^. Codon-optimized CAR19LH was cloned into third-generation lentiviral vector backbone pCCL-c-UBC (ubiquitin C promoter). The eGFP fluorescent protein coding sequence was added downstream of CD3ζ, separated by furin-2A linker for polycistronic expression.

Packaging and concentration of lentivirus particles were performed as previously described^19^. Briefly, 293T cells (Cat:CRL-3216; ATCC) were co-transfected with lentiviral vector plasmids pCMV-DR8.9 and pCAGGS-VSVG using TransIT-293 (Cat: MIR2704, Mirus BIO). After 17 h, cells were treated with 20 mM sodium butyrate (Cat: A11079.36, Thermo Fisher Scientific) for 8 h, followed by 48 h incubation in UltraCULTURE™ serum-free media (Cat: 12-725F, Lonza). Supernatants were concentrated by ultrafiltration using Amicon Ultra-15 100K Centrifugal filters (Cat: UFC910024, Millipore Sigma) at 4,000*g* for 40 min at 4 °C and stored at -80 °C.

### Design and validation of CRISPR/Cas9 guide RNAs

Guide RNAs (gRNA) were designed using published algorithms, including Benchling web tool (https://benchling.com) and Synthego CRISPR Design Tool (https://synthego.com). 2-3 gRNAs were chosen based on optimal silico-predicted on-and off-target scores (out of 100) and proximity to the endogenous stop codon (< 0-100 bp). Cutting efficiency was assayed *in vitro* by electroporation of pre-complexed CRISPR/Cas9 ribonucleoprotein (RNP) to T-iPSCs. Guides targeting exon 1 of *TRAC* and *B2M* were used as previously described^16,22^; newly designed sgRNA sequences were as follows: *GZMA* (5’-GAAGGTGTTTCATTACAGCG-3’), *CD8β* (5’- GTCCTGCTACAAAAAGACAT-3’), *TRAC* (5’-AGAGTCTCTCAGCTGGTACA-3’). Genomic DNA was collected from electroporated cells and the regions flanking the cleavage sites were amplified by PCR. Purified PCR products were sequenced using Sanger sequencing and on-target cutting efficiency was measured using the Tracking of Indels by Decomposition (TIDE) tool, which calculates the percentage of insertions or deletions (indels).

### CRISPR/Cas9 editing of T-iPSC lines

Genetic engineering of T-iPSCs was performed by delivering pre-complexed CRISPR/Cas9 ribonucleoprotein (RNP) using electroporation. Briefly, RNP complexation was achieved by incubating 90 pmol of custom-designed sgRNA (Synthego) and 60 pmol of purified spCas9 protein (QB3 MacroLab, University of California Berkeley) for 15 minutes at room temperature. Double-stranded plasmid HDR template was designed as follows: 5’ homology arm (∼1000 bp amplified by PCR) flanking the gRNA targeting sequence, furin-GSG-P2A self-cleaving peptide in frame with the last exon of the endogenous gene, followed by the *CAR19BB, CAR19LH* or *CAR20* cDNA. Anti-CD19 CAR19LH (scFv: FMC63) and anti-CD19 CAR19BB (scFv: FMC63) were designed and generated as previously described^7^. Anti-CD20 CAR20 features Leu-16 scFV, IgG4 hinge, CH2-CH3 (L235E, N297Q), CD28 transmembrane domain, CD28 co-stimulatory domain, and CD3ζ intracellular signaling domain. The donor template also included a *loxp* flanked puromycin resistance gene, followed by the 3’ homology arm (∼1000 bp amplified by PCR). Following incubation, 2 µg of the HDR template was combined with complexed RNP. Knockout of *B2M* was achieved as previously described^16^.

For electroporation, T-iPSCs cultured on Matrigel-coated plates were dissociated upon reaching 60% confluency using TrypLE™ Express (Thermo Fisher Scientific) and resuspended at a concentration of 2.5 x 10^5^ cells in 20 μl of pre-mixed P3 Primary Cell 4D-Nucleofector solution (P3 Primary Cell 4D-Nucleofector X Kit S, Cat: V4XP-3032; Lonza). Cells were combined with the RNP/HDR plasmid solution and transferred into individual wells of a 16-well Nucelocuvette Strip (Lonza). Nucleofection was performed using 4D-Nucleofector Core (Cat: AAF1003B, Lonza) and X Unit (Cat: AAF-1003X, Lonza) with the pulse code CB-150. Following electroporation, cells were rested at room temperature for 10 minutes and transferred to a well of a Matrigel-coated 12-well plate with 1 ml of mTeSR™ with Y-27632 dihydrochloride (10 uM). Cell culture medium was changed daily with Y-27632 dihydrochloride (10 uM) for 48-72 hours and passaged for puromycin selection.

### Puromycin selection and single-cell cloning of edited human T-iPSC lines

Puromycin selection of edited T-iPSCs was used to enrich for successfully edited cells. Briefly, edited T-iPSCs were dissociated into single-cell solution using TrypLE™ Express (Thermo Fisher Scientific) and plated at a density of 1.0 x 10^5^ cells/well of a Matrigel-coated 6-well plate in mTeSR™ and Y-27632 dihydrochloride (10 μM). After 24 hours, cell medium was changed daily with mTeSR™ supplemented with 0.50 ug/ml puromycin (Cat: 73342, Stem Cell Technologies) until cells reached 80% confluency.

Single-cell cloning of puromycin-selected T-iPSCs was achieved with low-density plating. Following puromycin selection, T-iPSCs were dissociated into single-cell solution with TrypLE™ Express and plated at a cell density of 2.5 x 10^3^ cells/plate of a Matrigel-coated 100 mm culture dish in mTeSR™ and with Y-27632 dihydrochloride (10 μM), which was removed after 24 h. Cell culture medium was changed daily. After colony formation, 12-24 individual colonies were scraped with a P200 pipette tip under a microscope and plated into individual wells of a Matrigel-coated 12-well plate with mTeSR™. Once colonies reached ∼70% confluency, T-iPSCs were dissociated using TrypLE™ Express and genomic DNA was isolated for genotyping to determine mono- or biallelic insertion of transgene.

### Lentiviral transduction of human T-iPSC lines

CAR19LH-transduced T-iPSC line was generated by lentiviral transduction of unedited T-iPSCs using a lentiviral vector encoding for CAR19LH-2A-eGFP. Briefly, T-iPSCs were dissociated into single-cell solution using TrypLE™ Express (Thermo Fisher Scientific) and plated at a density of 1.0 x 10^5^ cells/well of a Matrigel-coated 6-well plate in mTeSR™ and Y-27632 dihydrochloride (10 μM). After 24 hours, culture medium was replaced with 1 ml of mTeSR Plus and concentrated lentiviral supernatant was added directly into the well. Transduced T-iPSCs were cultured for 10-14 days and highest 5% eGFP+ T-iPSCs were isolated via FACS.

### Generation of human embryonic mesodermal progenitors

Mesodermal commitment was induced as previously described^41^. Briefly, expanded T-iPSCs were dissociated into single-cell solution using TrypLE™ Express (Thermo Fisher Scientific) and resuspended at a concentration of 1 x 10^6^ cells/ml in X-VIVO™ 15 Hematopoietic Serum-free Culture Media (Cat: BEBPO4-744Q, Lonza) supplemented with recombinant human (rh)activin A (10 ng ml^-1^, Cat: 338-AC-010, R&D Systems), rhBMP4 (10 ng ml^-1^, Cat: 314-BP-010, R&D Systems), rhVEGF (10 ng ml^-1^, Cat: 298-VS-005, R&D Systems), rhFGF (10 ng ml^-1^, Cat: 233-FB-025, R&D Systems), and Y-27632 dihydrochloride (10 uM). 3.0 x 10^6^ cells were plated in a well of a Matrigel-coated 6-well plate and medium was changed daily with X-VIVO™ 15 supplemented with rhBMP4 (10 ng ml^-1^), rhVEGF (10 ng ml^-1^), and rhFGF (10 ng ml^-1^). After 3.5 days, cells were dissociated into single-cell solution using Accutase (Cat: AT104, Innovative Cell Technologies Inc.) and resuspended in magnetic-activated cell soring (MACS) buffer (PBS, 2 mM EDTA, 0.5% bovine serum albumin) for flow cytometry analysis to evaluate differentiation with anti-CD56 and anti-CD326 prior to EMO generation.

### Generation of human embryonic mesodermal organoids

Embryonic mesodermal organoids (EMOs) were generated as previously described^16^. Briefly, EMOs were generated by the aggregation of EMPs with MS5-hDLL4 cells. MS5-hDLL4 cells were collected upon reaching ∼80% confluency using Trypsin-EDTA (Cat: 25300120, Thermo Fisher Scientific). MS5-hDLL4 and hEMPs were resuspended in EGM-2 Endothelial Cell Growth Medium-2 (Cat: CC-3162, Lonza) with Y-27632 dihydrochloride (10 μM) and SB-431542 (10 μM; Cat: 1614, Tocris Bioscience). 5 x 10^5^ MS5-hDLL4 cells were combined with 9 x 10^4^ EMP per EMO in a 1.7 ml Eppendorf tube (up to 144 EMOs per tube) and resuspended in EGM-2 media with Y-27632 dihydrochloride (10 μM) and SB-431542 (10 μM) at a volume of 6 μl per EMO. Three EMOs were plated individually (6 μl per EMO) on top of a 0.4 mm Millicell cell culture insert (Cat: PICM0RG50, Millipore) placed in a 6-well plate with 1 ml of EGM-2 media. EMO media was changed every 2 days with EGM-2 media and SB-431542 (10 uM). After 7 days, media was changed every 2 days with EGM-2 supplemented with SB-431542 (10 uM), rhTPO (5 ng ml^-1^, cat: 288-TPN-025, R&D Systems), rhFLT3L (5 ng ml^-1^, cat: 308-FK-025, R&D Systems), and rhSCF (50 ng ml^-1^, cat: 255-SC-200, R&D Systems) until day 14 of culture.

### Generation of artificial thymic organoids

Artificial thymic organoids (ATOs) were generated as previously described^16^. Briefly, EMOs were disaggregated using physical dissociation with a P1000 pipette tip and MACS buffer (PBS, 2 mM EDTA, 0.5% bovine serum albumin). After disaggregation, the cell solution was passed through a 70 μm nylon filter (Cat: 352350, Corning) and resuspended in ‘ATO culture media’, which consisted of RPMI 1640 (Cat: 10-040-CV, Corning), 4% B27 Supplement (Cat: 17504044, Thermo Fisher Scientific), 1% GlutaMAX Supplement (Cat: 35050061, Thermo Fisher Scientific), 30 mM L-Ascorbic acid 2-phosphate sesquimagnesium salt hydrate resuspended in PBS (Cat: A8960-5G, Sigma-Aldrich), 1% Penicillin-Streptomycin (Cat: 15140122, Thermo Fisher Scientific), supplemented with rhSCF (50 ng ml^-1^), rhFLT3L (5 ng ml^-1^), and rhIL-7 (5 ng ml^-1^, Cat: 207-IL-010, R&D Systems). 1 x 10^4^ post-EMO cells were combined with 2.5 x 10^5^ MS5-hDLL4 stromal cells per ATO in a 1.7 ml Eppendorf tube (up to 144 ATOs per tube) and resuspended in ‘ATO culture media’ at a volume of 6 μl per ATO. Three ATOs were plated on top of individual 0.4 mm Millicell cell culture insert (Millipore) with 1 ml media. Complete ‘ATO culture media’ was changed every 3-4 days for the duration of the experiment (5-7 weeks).

### Isolation of ATO-derived T cells

DP or SP8 T cells were isolated from ATO culture at week 2 or weeks 5, respectively. Briefly, ATOs were harvested by physical disaggregation from each filter using a P1000 pipette tip and MACS buffer (PBS, 2 mM EDTA, 0.5% bovine serum albumin) and passed through a 70 μm nylon strainer. Up to 90 ATOs were collected in a single well filled with MACS buffer before being transferred through the strainer. Aggregates were further dissociated on top of the strainer using the back end of a sterile 1 ml syringe. After collection, the filter was washed with MACS buffer and centrifuged at 300 x g for 5 min at 4°C in a swinging bucket centrifuge.

### Stimulation of ATO-derived T cells

SP8 or DP T cells were isolated as described above and stimulated *in vitro* using irradiated K562 CD19-CD137L aAPCs at a 1:3 aAPC:T cell ratio in X-VIVO™ 15 (Lonza), supplemented with 5% human AB serum (Cat: 100-512-100, Gemini Bio), rhIL-2 (20 ng ml^-1^, Cat: 200-02, Peprotech) and rhIL-7 (5 ng ml^-1^). Fresh T cell media was replenished every 2-3 days and replated into larger wells as necessary. Re-stimulations were performed every 7 days. Prior to assays, T cells were rested for a minimum of 3-5 days in media without cytokines at the end of the stimulation cycle.

### T cell activation assay

SP8 T cells were isolated from ATO cultures at week 5 as described above and were rested for 3-5 days (Tn-like) or underwent two stimulation cycles (Tem-like). 2.5 x 10^5^ SP8 T cells were co-cultured with irradiated K562 cells expressing cognate antigen (CD19-K562) or no antigen (K562) at a 1:1 Target:T cell ratio in 96-well U-bottom plates with 250 µl X-VIVO™ 15 (Lonza) supplemented with 5% human AB serum (Gemini Bio) for 24 h at 37 °C. Cells were then washed and stained for CD3, CD7, CD4, CD8a, CD25, and CD137 (Biolegend). DAPI was added immediately prior to flow cytometry analysis.

### T cell cytokine release assay

SP8 T cells were isolated and stimulated or rested as described above. 2.5 x 10^5^ SP8 T cells were co-cultured with irradiated K562 target cells expressing cognate antigen (CD19-K562) or no antigen (K562) at a 1:1 Target:T cell ratio in a 96-well U-bottom plates and 250 µl X-VIVO™ 15 (Lonza) supplemented with 5% human AB serum (Gemini Bio) and human Protein Transport Inhibitor Cocktail (1X, Cat: 00-4980-03, eBioscience) for 6 h at 37 °C. Following 4 h of incubation, CD107a-APC antibody (Biolegend) was added to each well at a 1:10 dilution. Cells were washed and stained for CD3, CD4, CD7, CD8a, and Zombie UV Fixable Viability Dye (Biolegend) prior to fixation and permeabilization using an Intracellular Fixation & Permeabilization Buffer (Cat: 88-8824-00, eBioscience) and intracellular staining for IFNγ and TNFα (Biolegend).

### Incucyte cytotoxicity and rechallenge assays

SP8 T cells were isolated and expanded or rested as described above. Each well of a 96-well Clear Bottom Microplate (Cat: 3904, Corning) was first coated with 50 μl of poly-L-lysine solution (Cat: P4797, Sigma Aldrich) for 1 hour at room temperature, followed by 3 washes with 200 μl PBS. 1 x 10^4^ SP8 T cells were co-cultured with K562 target cells expressing cognate antigen (CD19-K562-mKate2) or no antigen (K562-mKate2) at a 1:1 Target:T cell ratio in 250 μl X-VIVO 15 media (Cat: 02-053Q, Lonza), supplemented with 5% human AB serum (Gemini Bio), IL-2 (20 ng ml^-1^), and rhIL-7 (5 ng ml^-1^). Triplicate wells were prepared for each condition and live cell imaging was performed every 2 h on Incucyte SX5 (Sartorius) for 120 h. Red fluorescence was evaluated to measure the growth of target cells at each time point using Incucyte Software (v2020C).

For repeated tumor challenge experiments, T cell co-culture conditions were first prepared as described above using 1.5 x 10^4^ SP8 T cells that were co-cultured with K562 target cells expressing cognate antigen (CD19-K562-mKate2) or no antigen (K562-mKate2) at a 1:1 Target:T cell ratio. 1.5 x 10^4^ K562 target cells were added to each well every 72 h until the end of 4 total challenge cycles. T cell media was replenished during each rechallenge period. Red fluorescence was evaluated to measure the growth of target cells every 2 h during the course of the experiment using Incucyte SX5 (Sartorius).

### Single-cell 10X multiome library preparation and sequencing

The nuclei of ATO-derived T cells were isolated and submitted for 10X Multiome library prep. Briefly, at 3 weeks of ATO culture, total ATO-derived cells were isolated as described above using PBS + 0.04% BSA (GeminiBio). Isolated cells were first processed using MACS Dead Cell Removal Kit (Cat: 130-090-101, Miltenyi Biotec) and 1 x 10^6^ cells were transferred to a 1.7 ml Eppendorf tube. Cells were spun down and lysed for 4 min using chilled lysis buffer that consisted of nuclease-free water (Cat: J71786.AE, Thermo Fisher), Tris-HCL (10 mM, Cat: T2194, Sigma-Aldrich), NaCl (10 mM, Cat: 59222C, Sigma-Aldrich), MgCl_2_ (3 mM, Cat: M1028, Sigma-Aldrich), BSA (1%), Tween-20 (0.1%, Cat: 1662404, Bio-Rad), IGEPAL CA-630 (0.1%, Cat: i8896, Millipore Sigma), Digitonin (0.01%, Cat: BN2006, Thermo Fisher), DTT (1 mM, Cat: 646563, Sigma-Aldrich), and RNase inhibitor (1 U μl^-1^, Cat: 3335402001, Sigma-Aldrich). Cell lysis was spun down and washed 3 times with chilled wash buffer, which consisted of nuclease-free water, Tris-HCL (10 mM), NaCl (10 mM), MgCl_2_ (3 mM), BSA (1%), Tween-20 (0.1%) DTT (1 mM), and RNase inhibitor (1 U μl^-1^). Isolated nuclei were spun down at 500 g for 5 min and resuspended with chilled Nuclei Buffer, which consisted of nuclease-free water, 10X Nuclei Buffer (1X, Cat: PN-1000284, 10X Genomics), DTT (1 mM), and RNase inhibitor (1 U μl^-1^) at a concentration of 5-8 x 10^3^ nuclei μl^-1^. Nuclei were provided to the Technology Center for Genomics and Bioinformatics (TCGB) core at UCLA for Chromium Next GEM Single Cell Multiome library prep (10X Genomics). Fully constructed libraries for all samples were sequenced on a 10B flow cell on the Illumina NovaSeq X Plus System.

### Multiome sequencing data filtration and cleaning

Sequenced reads from each sample were aligned to the human reference genome GRCh38 (cellranger-arc-GRCh38-2020-A-2.0.0, 10X Genomics) and processed using the Cell Ranger ARC v.2.0.2 (10X Genomics) “count” pipeline to generate multiomic count matrices for both GEX and ATAC libraries. We achieved a minimum of ∼24,000 mean reads per cell for GEX libraries, and >30,000 mean reads per cell for ATAC libraries.

GEX and ATAC count matrices from each sample were analyzed with the Seurat v.4.4.0 (Satija Lab) and Signac v.1.13 packages in R v.4.3.2^42^. Briefly, Seurat objects were created from the output matrices from the Cell Ranger ARC pipeline including both GEX and ATAC counts as well as fragments.tsv.gz, which was annotated using EnsDb.Hsapiens.v86. After loading, individual datasets were bioinformatically cleaned for doublets using DoubletFinder v.2.0.4^43^. Then, to ensure the use of high-quality cells, cleaned barcodes were filtered to remove cells with low UMI counts (nUMI ≥ 1000), low number of sequenced genes (nGene ≥ 1000), high mitochondrial gene expression due to cellular stress or loss of RNA (mitoRatio < 0.18). Additionally, the dataset was cleaned for cells expressing human DLL4 as we have found multilineage clustering in unsorted samples, perhaps due to myeloid cell interaction with MS5-hDLL4 stromal cells in the organoid^18^.

After initial data filtration for doublets, low-quality, and outlier cells, the combined Seurat object was split by each modality, RNA and ATAC, and then batch corrected for technical and biological variations using the Reciprocal Principal Component Analysis (RPCA) integration method in Seurat and the reciprocal LSI projection method in Signac, respectively^42,44^. For integration of the combined RNA modality, molecular count data for each sample were individually normalized and variance stabilized using SCTransform, which bypasses the need for pseudocount addition and log-transformation, and then cell cycle phase scores were calculated for each individual sample based on the expression of canonical cell cycle genes within a specific barcoded cell. Following cell cycle scoring, raw counts were normalized, and variance stabilized again using SCTransform with the additional steps of regressing mitochondrial gene expression and calculated cell cycle scores in order to mitigate their effects on downstream projections due to the heterogeneity of *in vitro* organoid cultures. In order to perform RPCA integration, highly variable genes (nfeatures = 5000) were then identified from each sample and then used to find integration anchors between datasets (k.anchor = 10). Integration of the ATAC modality was performed as detailed in the Signac integration vignette^44^. The uncorrected low-dimensional cell embeddings (LSI) were calculated using the merged dataset, and integration was performed by identifying anchors between the individual samples by projecting them into the shared low-dimensional space that was calculated. Visualization of aggregated signal from genomic regions with genome annotations was performed with the CoveragePlot() function in Signac^44^.

### Joint visualization and analysis of scRNA-seq and scATAC-seq using the weighted nearest neighbor method

Integrated scRNA-seq and scATAC-seq datasets from all samples were clustered and visualized using the weighted nearest neighbor method (WNN) in Seurat and the IKAP package^42,45^. IKAP identified the optimal principal components (PC=13) for identifying multimodal neighbors from the RNA modality and WNN UMAP projection and determined the individual clustering of our populations. Clusters were identified via markers identified through the FindAllMarkers() function in Seurat, and DP and SP8 clusters were further subset using additional iterations of IKAP to identify DP-1, DP-2, Intermediate SP8, SP8-1, and SP8-2 clusters. Notably, one cluster (NK), separated from the main body of cells in the dimensionally reduced WNN UMAP projection, was removed from the dataset based on irregular, multilineage gene expression.

Differentially expressed gene analysis of scRNA-seq datasets

Given the individual SCTransform models calculated from each sample, the minimum median UMI was used to recorrect the counts and data slot for downstream analysis using PrepSCTFindMarkers in Seurat (Satija Lab)^42^. Differentially expressed genes (DEGs) were calculated using the “MAST” algorithm72 which is tailored to scRNA-seq data DEG analysis using a model that parameterizes both stochastic dropout and characteristic bimodal expression distributions, for the FindMarkers function of Seurat (min.pct = 0.25, logfc.threshold = 0.25). DEGs from FindMarkers were used to generate ranked gene lists ordered by log-fold change for Gene Set Enrichment Analysis (GSEA) using the fgsea v1.22.073 package and gene signatures were pulled from the Molecular Signatures Database (MSigDB) using msigdbr v7.15.174 (<https://CRAN.R-project.org/package=msigdbr>). Visualization of GSEA results was performed using the enrichplot v1.16.2 package75 (<https://yulab-smu.top/biomedical-knowledge-mining-book/>). Additional gene enrichment pathway analysis and gene annotations using DEGs were performed using Metascape^24^ (<https://metascape.org/gp/index.html#/main/step1>).

### Trajectory inference and pseudotime analysis

The integrated dataset from before was subset for the main T cell developmental populations and then converted to a CellDataSet using the as.cell_data_set() function from SeuratWrappers. Trajectory inference was performed using monocle3 v.1.3.7 and the WNN UMAP reduction to reconstruct the developmental trajectories of the cells^46^. Pseudotime values were calculated by identifying the earliest node found within the “DP” clusters and selecting it as the root node of the trajectory using the interactive order_cells function in monocle3. Calculated pseudotime trajectories were visualized using the WNN UMAP with cell colors corresponding to calculated pseudotime values. In order to visualize gene expression changes over pseudotime, pseudotime values and gene expression matrices were extracted and visualized using the ggplot2 package (<https://ggplot2.tidyverse.org>).

### Flow cytometry analysis and antibodies

All staining for flow cytometry analysis and sorting was performed in MACS buffer for 20 minutes at 4 °C in the dark. TruStain FcX (Cat: 422302, Biolegend) was added to all samples for 5-10 minutes prior to antibody staining. All anti-CD19 CARs were detected by first performing a primary incubation using biotinylated anti-FMC63 antibody (Cat: FM3-BY54-25tests, ACROBiosystems) followed by a secondary incubation using Biotin-PE antibody (Biolegend). DAPI was added to all sample immediately prior to analysis.

Flow cytometry analysis was performed on BD LSRII Fortessa, and FACS sorting was performed on BD FACSAria instrument (BD Biosciences) with the assistance of the UCLA Broad Stem Cell Research Center Flow cytometry Core. All BD instruments used BDFACSDiva v8.0.2 software (BD Biosciences). Flow cytometry data were analyzed using FlowJo v10.10 software (BD Biosciences). For all flow cytometry analysis, doublets were initially removed through forward scatter (FSC) height versus FSC-width and side scatter (SSC) height versus SSC-width gating and DAPI/Zombie-positive cells were gated out.

Human antibody clones used in these studies were obtained from Biolegend: CD3 (UCHT1), CD4 (RPA-T4), CD5 (UCHT2), CD7 (CD7-6B7), CD8α (SK1), CD19 (HIB19), CD20 (2H7), CD25 (BC96), CD27 (O323), CD28 (CD28.2), CD45 (HI30), CD45RO (UCHL1), CD45RA (HI100), CD56 (5.1H11), CD62L (DREG-56), CD107a (H4A3), CD197 (G043H7), TCRαβ (IP26), Biotin (1D4-C5), B2M (2M2), TNF-α (MAb11), IL-2 (MQ1-17H12), IFN-γ (4S.B3), CD366 (F38-2E2), CD223 (11C3C65), CD152 (L3D10), TIGIT (A15153G), CD279 (EH12.2H7). The anti-human antibody CD8β (REA715) was obtained from Miltenyi Biotec. The anti-mouse antibody mCD29 (HMβ1-1) was obtained from Biolegend. The antibody against the CD19 CAR (FMC-63) was obtained from ACROBiosystems.

### Statistics

In all figures, data are presented as mean ± standard deviation (SD) as indicated. Exact *n* values represent independent experiments and are specified in each figure legend. Statistical tests were conducted using GraphPad Prism software using the two-tailed unpaired *t-*test and ordinary one-way ANOVA. *P* value significance was indicated directly on each figure and classified as such: \**p* < 0.05, \*\**p* < 0.01, \*\*\**p* < 0.001, \*\*\*\**p* < 0.0001.

## Data availability

The main data supporting the results in this study are available within the paper and its Supplementary information. The genomic data used in Fig. 1b and 2a were extracted from GSE116015^19^ and PRJNA741323^20^. The raw and analyzed datasets generated during this study for gene expression from this study will be available through the NCBI Gene Expression Omnibus repository upon acceptance for publication. The data that support the findings of this study are available from the corresponding author upon request.

## Reporting Summary

Further information on research design is available in the Nature Research Reporting Summary linked to this article.

## Acknowledgements

We thank J. Scholes, F. Codrea and J. Calimlim from the UCLA Broad Stem Cell Research Center (BSCRC) Flow Cytometry core, the TCGB Shared Resource of the NCI-designated UCLA Jonsson Comprehensive Cancer Center (NIH/NCI [P30CA016042]), Y. Chen, H. Mikkola, and M. Su for their collective guidance on stem and T cell biology, R. Crisostomo for assistance with gene editing, and A. Montel-Hagen for optimization of the ATO culture system. The research presented in this article was supported in part by NIH/NCI [F30CA278297] (S.P.Y), NIH/NCI [F31CA239555] (P.C), UCLA BSCRC fellowships (S.P.Y. and X.Y.), UCLA-Caltech Medical Scientist Training Program NIGMS [T32GM008042] (S.P.Y), NIH/NHLBI T32HL086345 (P.C), NIH/NCI [K08CA235525] (C.S.S.), V Foundation Scholar Award (C.S.S.), Parker Institute of Cancer Immunotherapy (C.S.S.), and a sponsored research agreement between UCLA and Pluto Immunotherapeutics, Inc (G.M.C). This research was also made possible in part by grants to G.M.C. from the California Institute for Regenerative Medicine [GC1R-06673-B: CRP and DISC2-10134]. The contents of this publication are solely the responsibility of the authors and do not necessarily represent the official views of CIRM or any other agency of the State of California.

## Author contributions

S.P.Y conceptualized and performed all experiments, analysed data, prepare figures, and wrote and edited the article. X.Y. assisted with the incucyte *in vitro* experiments and edited the article. C.E. and S.S. assisted with gene editing and cloning of T-iPSC lines. P.C.C. assisted with sequencing data processing and analysis. L.L., J.L., and A.A. assisted with stem cell and ATO cultures. V.C. and H.M. helped spearhead the initial sequencing and analysis of bulk RNA-sequencing data. S.L., C.S.S., and D.B.K. provided critical materials and conceptual advice and guidance. G.M.C. directed the project and co-wrote and edited the article.

