## Supplemental Figures for "Stage-specific CAR-mediated signaling generates naïve-like, TCR-null CAR T cells from induced pluripotent stem cells"

### Supplementary Figures

#### Supplementary Figure 1: Generation of *GZMA*-mCitrine reporter T-iPSC and analysis of ATO development

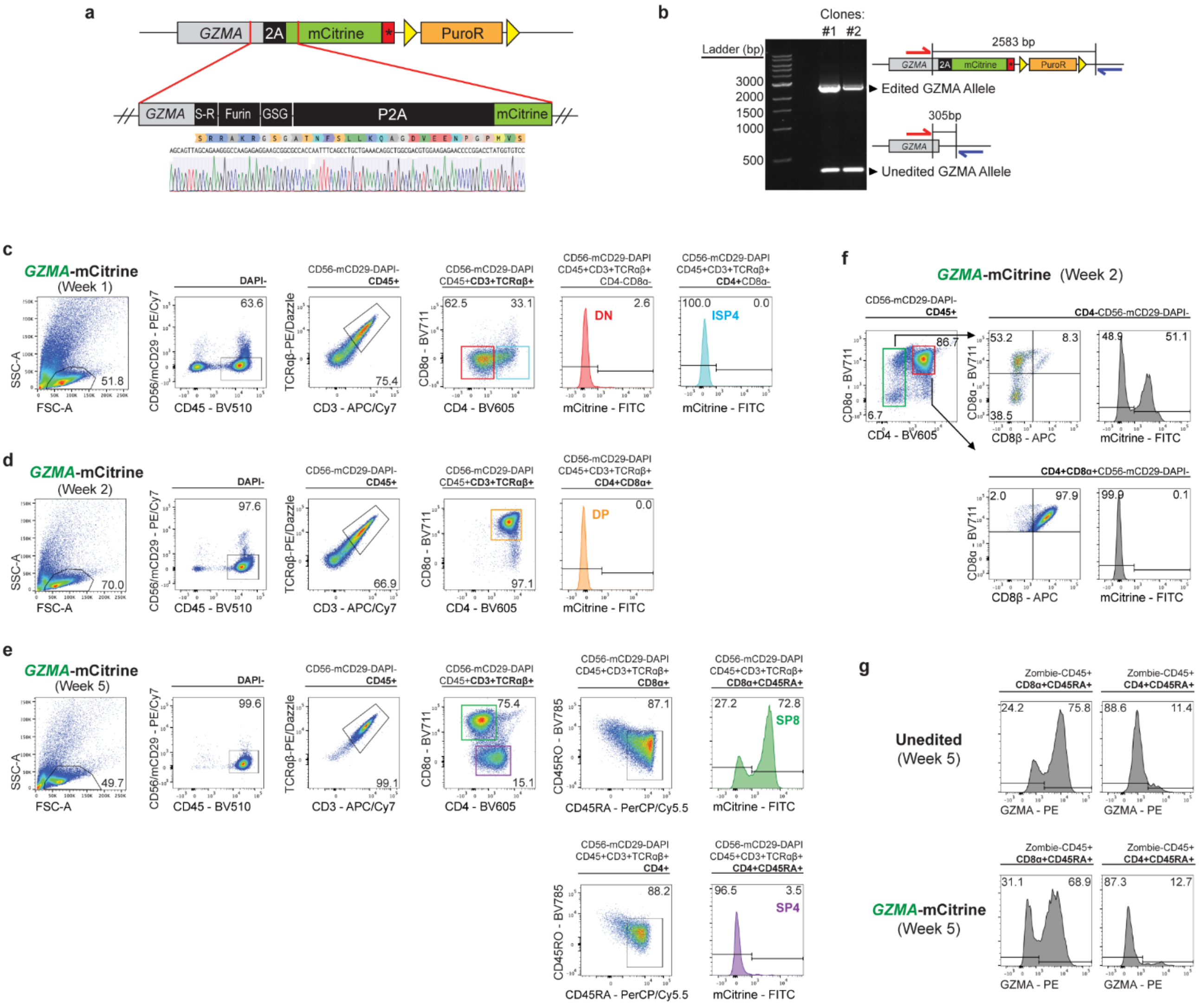

**Supplementary Figure 1: Generation of *GZMA*-mCitrine reporter T-iPSC and analysis of ATO development**

**(a)** Sanger sequencing of *GZMA* 3'UTR allele at the region of mCitrine transgene insertion. The last exon of *GZMA* was linked to the mCitrine transgene using a S-R linker (Ser-Arg), furin cleavable linker (furin), GSG linker (Gly-Ser-Gly), and P2A ribosome skip sequence.

**(b)** Gel electrophoresis of PCR products generated from the amplification of the region flanking *GZMA* sgRNA cut site. The size of the PCR product helped determine the insertion of the mCitrine transgene; the PCR product of the edited *GZMA* allele was 2583 bp and the unedited allele was 305 bp. The presence of the two bands in clones #1 and #2 indicates mCitrine transgene was successfully inserted in one allele of *GZMA*.

**(c-e)** Representative flow cytometry plots of the gating strategy used to measure mCitrine reporter expression at each stage of T cell development during *GZMA*-mCitrine ATO culture, evaluated at **(c)** 1, **(d)** 2 and **(e)** 5 weeks.

**(f)** Analysis of reporter expression in T cell precursors at week 2 of *GZMA*-mCitrine ATO culture. DN T cells gave rise to a transient population of mCitrine+ CD8αα+ T cells that expressed *GZMA*.

**(g)** Intracellular *GZMA* expression in SP8 and SP4 T cells generated from unedited (top) and *GZMA*-mCitrine T-iPSCs (bottom) at week 5 of ATO culture.

### Supplementary Figure 2: Analysis of *GZMA*-CAR19LH T-iPSC differentiation in ATOs

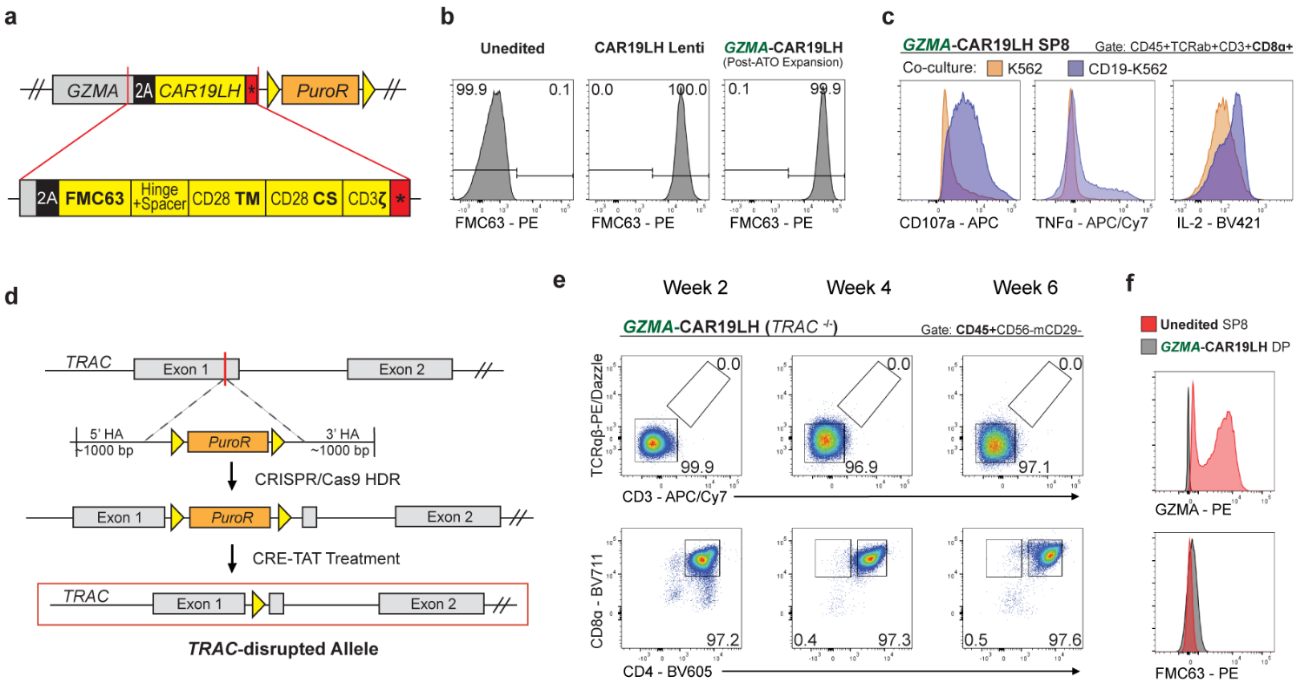


**Supplementary Figure 2: Analysis of *GZMA*-CAR19LH T-iPSC differentiation in ATOs**

**(a)** Schematic of second-generation CAR19LH (1928z) inserted into the *GZMA* locus. CAR19LH features an anti-CD19 scFv (clone: FMC63), IgG4 hinge with a CH2-CH3 spacer, CD28 transmembrane (TM) domain, CD28 co-stimulatory (CS) domain, and CD3ζ intracellular signaling domain.

**(b)** CAR19LH expression (FMC63) in total cells isolated from unedited, CAR19LH Lenti, and *GZMA*-CAR19LH ATOs. Total cells isolated from unedited and CAR19LH Lenti ATOs were analysed at week 5 of ATO culture. Total cells from *GZMA*-CAR19LH ATOs were first isolated at week 5 of ATO culture and underwent aAPC-mediated expansion prior to analysis. All cells were gated CD45+CD56-mCD29- (data representative of n = 5 independent experiments).

**(c)** CD107a and cytokine production (TNF-α, IL-2) in *GZMA*-CAR19LH SP8 T cells in response to 6h co-culture with CD19-K562 (purple) or K562 (orange) (data representative of n = 4 independent experiments).

**(d**) Schematic of gene editing strategy used to disrupt the *TRAC* locus. CRISPR/Cas9 HDR donor template was designed to insert a *loxp*-flanked (yellow triangle) puromycin resistance gene inside exon 1 of *TRAC*. Successfully edited iPSC clones underwent CRE-TAT treatment and a single *loxp* sequence remained, which interrupted the expression of *TRAC*. Clones with bi-allelic *TRAC*-disruption were used for ATO differentiation.

**(e)** Analysis of T cell development and endogenous TCR expression during *GZMA*-CAR19LH (*TRAC*$^{-/-}$) T-iPSC differentiation in ATOs at indicated time points (data representative of n = 3 independent experiments).

**(f)** Intracellular GZMA and surface CAR19LH (FMC63) expression in SP8 T cells generated from unedited T-iPSCs and DP T cells generated from *GZMA*-CAR19LH (*TRAC*$^{-/-}$) T-iPSCs at week 5 of ATO culture. SP8 T cells were gated CD45+CD8α+CD4-Zombie- and DP T cells were gated CD45+CD8α+CD4+Zombie- (data representative of n = 3 independent experiments).

### Supplementary Figure 3: Analysis of *CD8β*-CAR19BB T-iPSC differentiation in ATOs

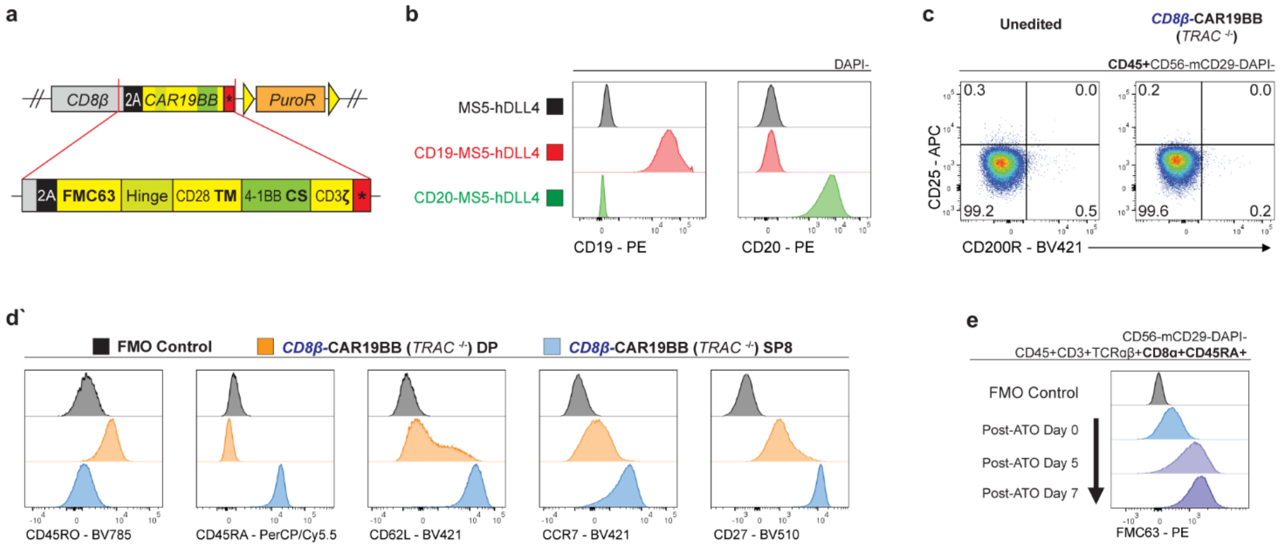


**Supplementary Figure 3: Analysis of *CD8β*-CAR19BB T-iPSC differentiation in ATOs**
**(a**) Schematic of second-generation CAR19BB (19BBz) inserted into *CD8β* locus. CAR19BB features an anti-CD19 scFv (clone: FMC63), IgG4 hinge, CD28 transmembrane domain, 4-1BB co-stimulatory domain, and CD3ζ intracellular signaling domain.

**(b)** Expression of CD19 or CD20 in engineered MS5-hDLL4 stroma lines used to generate ATOs.

**(c)** Expression of ILC2 markers CD25 and CD200R in unedited (left) and *CD8B*-CAR19BB (*TRAC*$^{-/-}$) (right) T cells analyzed at week 2 of ATO culture (data representative of n = 3 independent experiments).

**(d)** Phenotypic comparisons of *CD8B*-CAR19BB (*TRAC*$^{-/-}$) DP and SP8 T cells at week 6 of ATO culture compared to FMO controls. DP T cells were gated CD45+TCRαβ-CD3-CD8α+CD4+CD56-mCD29- and SP8 T cells were gated CD45+TCRαβ-CD3-CD8α+CD4-CD56-mCD29- (data representative of n = 6 independent experiments).

**(e)** CAR19BB expression (FMC63) in *CD8B*-CAR19BB (*TRAC*$^{-/-}$) SP8 T cells analyzed at week 6 of CD19-expressing ATO cultures (Post-ATO Day 0) and 5 and 7 days after isolation from ATOs (data representative of n = 3 independent experiments).

### Supplementary Figure 4: CAR-mediated positive selection was reproduced using different CARs and in the absence of Class I MHC

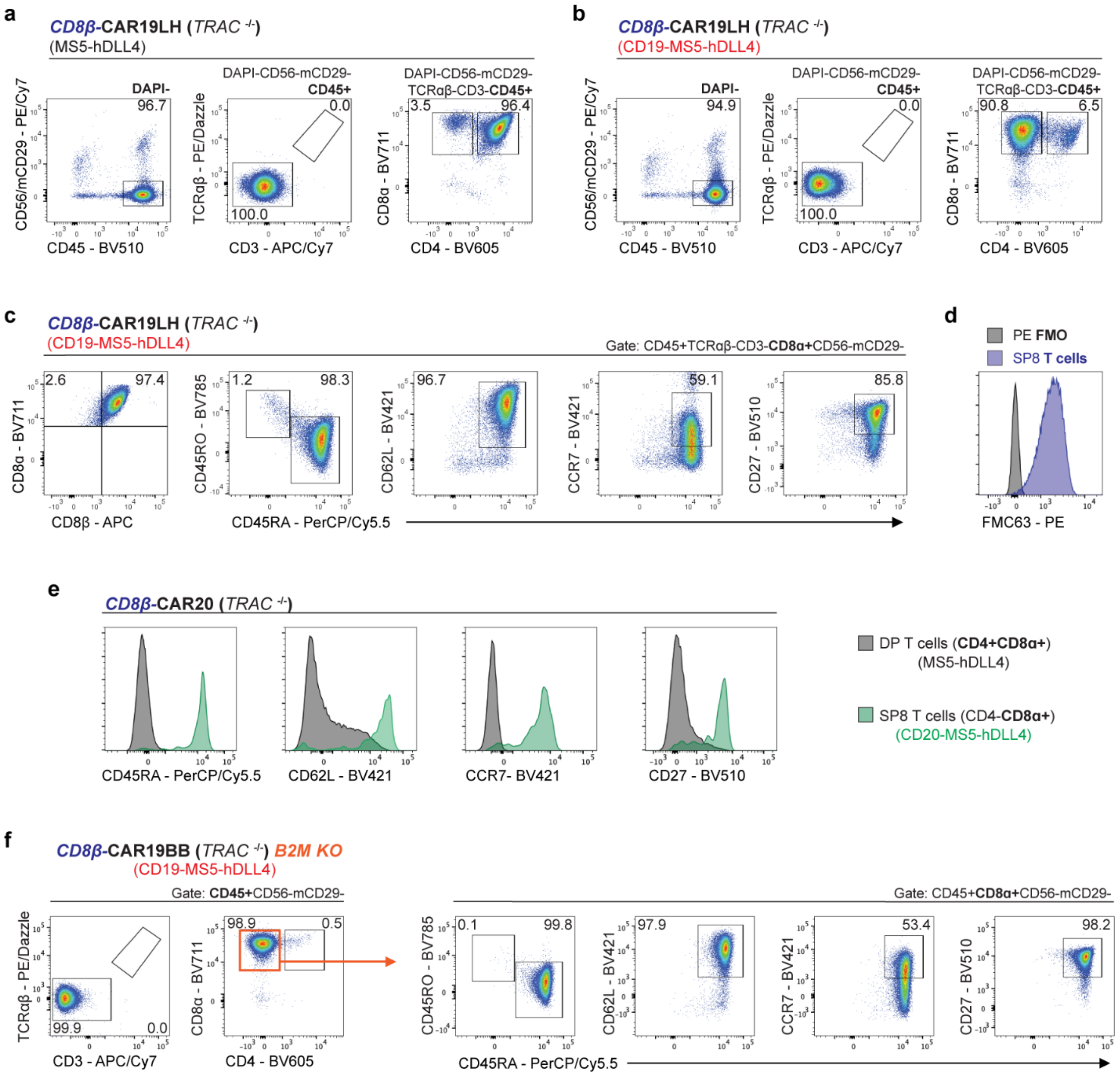

**Supplementary Figure 4: CAR-mediated positive selection was reproduced using different CARs and in the absence of Class I MHC**
**(a,b)** Analysis of T cell development in *CD8B*-CAR19LH (*TRAC*$^{-/-}$) ATOs generated using **(a)** MS5-hDLL4 or **(b)** CD19-MS5-hDLL4 at week 6 of ATO culture (data representative of n = 2 independent experiments).

**(c,d)** Analysis of **(c)** phenotype and **(d)** surface CAR19LH expression (FMC63) in *CD8B*-CAR19LH (*TRAC*$^{-/-}$) SP8 T cells analyzed at week 6 of ATO culture (data representative of n = 2 independent experiments).

**(e)** Phenotype of *CD8B*-CAR20 (2028z) (*TRAC*$^{-/-}$) T cells analyzed at week 4 of ATO culture in the presence or absence of CD20 antigen. DP T cells (CD45+TCRαβ-CD3-CD8α+CD4+CD56-mCD29-) are shown in black and SP8 T cells (CD45+TCRαβ-CD3-CD8α+CD56-mCD29-) are shown in green (data representative of n = 2 independent experiments).

**(f)** Analysis of T cell development in *CD8B*-CAR19BB (*TRAC*$^{-/-}$) *B2M* KO ATOs and SP8 phenotype at week 6 of culture (data representative of n = 2 independent experiments).

**Supplementary Figure 5: *CD8β*-CAR19BB SP8 phenotype during *ex vivo* stimulation**

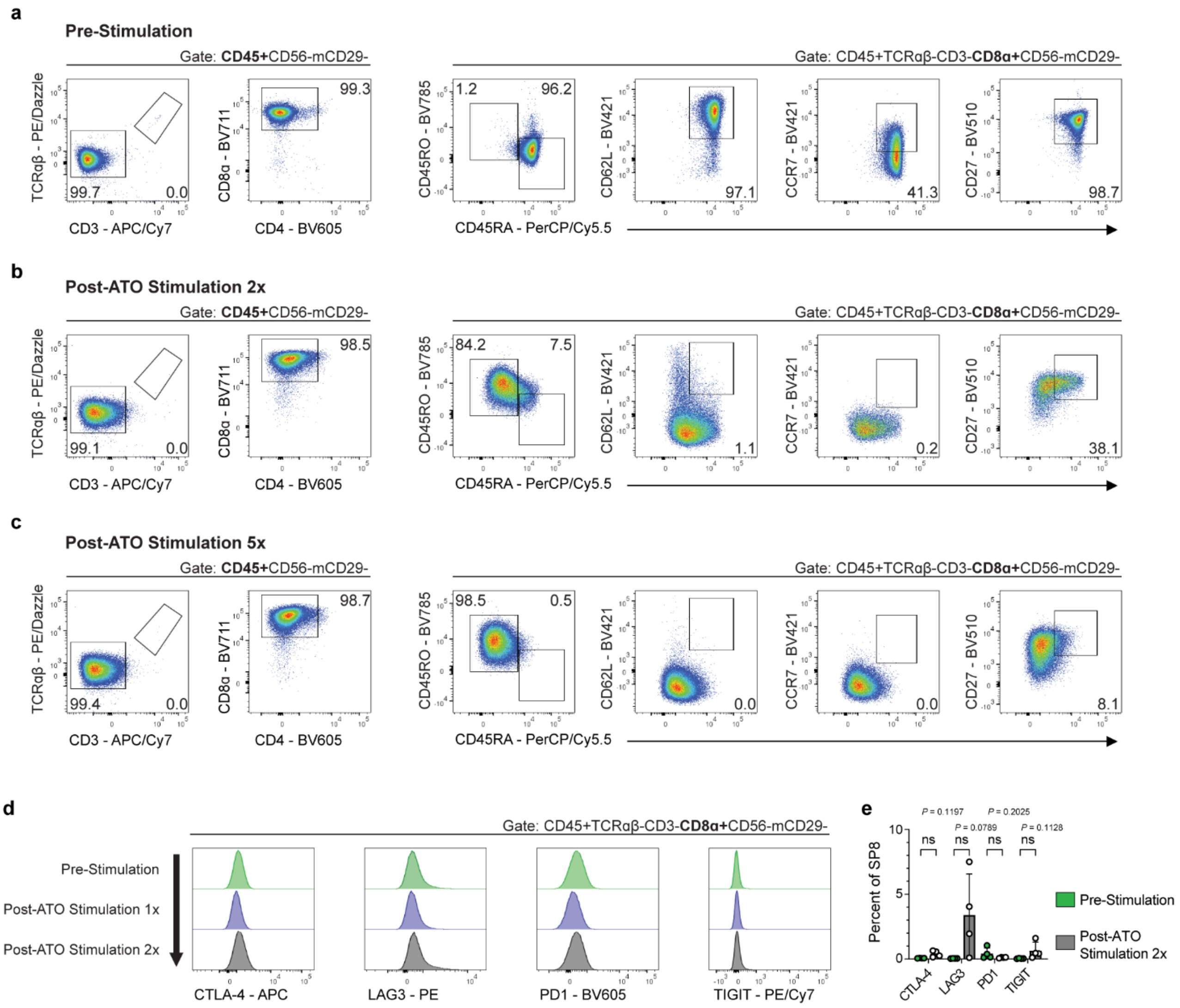


**Supplementary Figure 5: *CD8β*-CAR19BB SP8 phenotype during *ex vivo* stimulation**
**(a-c)** Phenotype of *CD8B*-CAR19BB (*TRAC*$^{-/-}$) SP8 analyzed at **(a)** week 5 of ATO culture (Pre-Stimulation) and after completing **(b)** 2 and **(c)** 5 cycles of aAPC-mediated stimulations using CD19-K562-CD137L (Post-ATO Stimulation) (data representative of n = 3 independent experiments).

**(d)** Expression of canonical exhaustion markers in *CD8B*-CAR19BB (*TRAC*$^{-/-}$) SP8 T cells analyzed at week 5 of ATO culture (Pre-Stimulation) and after 1 and 2 cycles of aAPC-mediated stimulation (Post-ATO Stimulation) (data representative of n = 3 independent experiments).

**(e)** Percent of *CD8B*-CAR19BB (*TRAC*$^{-/-}$) SP8 T cells expressing canonical exhaustion markers before (Pre-Stimulation) and after undergoing 2 cycles of aAPC-mediated stimulation (mean ± SD, **P* < 0.05 by multiple two-tailed unpaired *t*-tests, data representative of n = 3 independent experiments).

**Supplementary Figure 6: Functional characterization of Tn-like and Tem-like *CD8B*-CAR19BB (*TRAC*$^{-/-}$) SP8 T cells.**

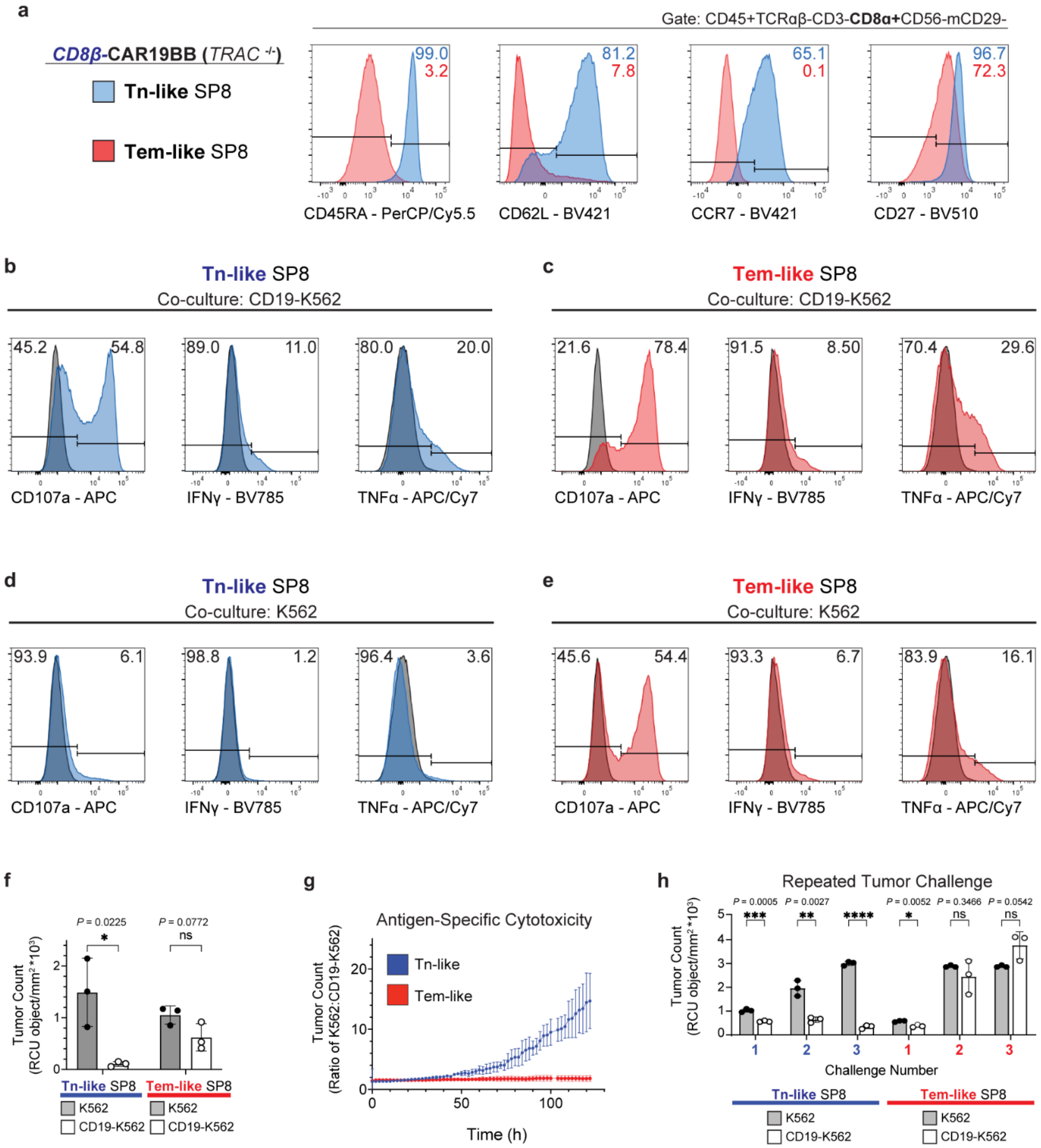

**Supplementary Figure 6: Functional characterization of Tn-like and Tem-like *CD8B*-CAR19BB (*TRAC*$^{-/-}$) SP8 T cells.**

**(a)** Phenotype of Tn-like and Tem-like *CD8B*-CAR19BB (*TRAC*$^{-/-}$) SP8 T cells. Tn-like SP8 T cells were isolated at week 5 of ATO culture (blue). Tem-like SP8 T cells were produced by first isolating SP8 T cells at week 5 of ATO culture and undergoing two 7-day stimulation cycles using CD19-K562-CD137L aAPCs (red). Unexpanded SP8 T cells retained a naïve phenotype, and expanded SP8 T cells differentiated into Tem-like cells.

**(b,c)** Representative histograms of CD107a expression and cytokine production (IFNγ, TNFα) in **(b)** Tn-like (blue) and **(c)** Tem-like (red) SP8 T cells in response to 6h co-culture with CD19-K562 compared to no co-culture controls (black). SP8 T cells were gated CD45+CD8α+Zombie- (data representative of n = 4 independent experiments).

**(d,e)** Representative histograms of CD107a expression and cytokine production (IFNγ, TNFα) in **(d)** Tn-like (blue) and **(e)** Tem-like (red) SP8 T cells in response to 6h co-culture with antigen negative K562 compared to no co-culture controls (black). SP8 T cells were gated CD45+CD8α+Zombie- (data representative of n = 4 independent experiments).

**(f)** Tumor growth measured at the final timepoint of 120h Incucyte cytotoxicity assay. Tn-like and Tem-like SP8 T cells were co-cultured with antigen negative K562 or CD19-K562 at a Target:T cell ratio of 1:1 (mean ± SD, **P* < 0.05 by multiple two-tailed unpaired *t*-tests, n = 3 independent experiments).

**(g)** Ratio of K562 to CD19-K562 tumor growth during a 120h Incucyte cytotoxicity assay. Higher ratio indicates higher antigen-specific clearance of tumor (mean ± SD, n = 3 independent experiments).

**(h)** Tumor growth measured at the end of each of the three re-challenge cycles using Incucyte assay (mean ± SD, *****P* < 0.0001 by multiple two-tailed unpaired *t*-tests, data representative of n = 2 independent experiments).

**Supplementary Figure 7: Development and phenotype of ATO cultures processed for single-cell sequencing**

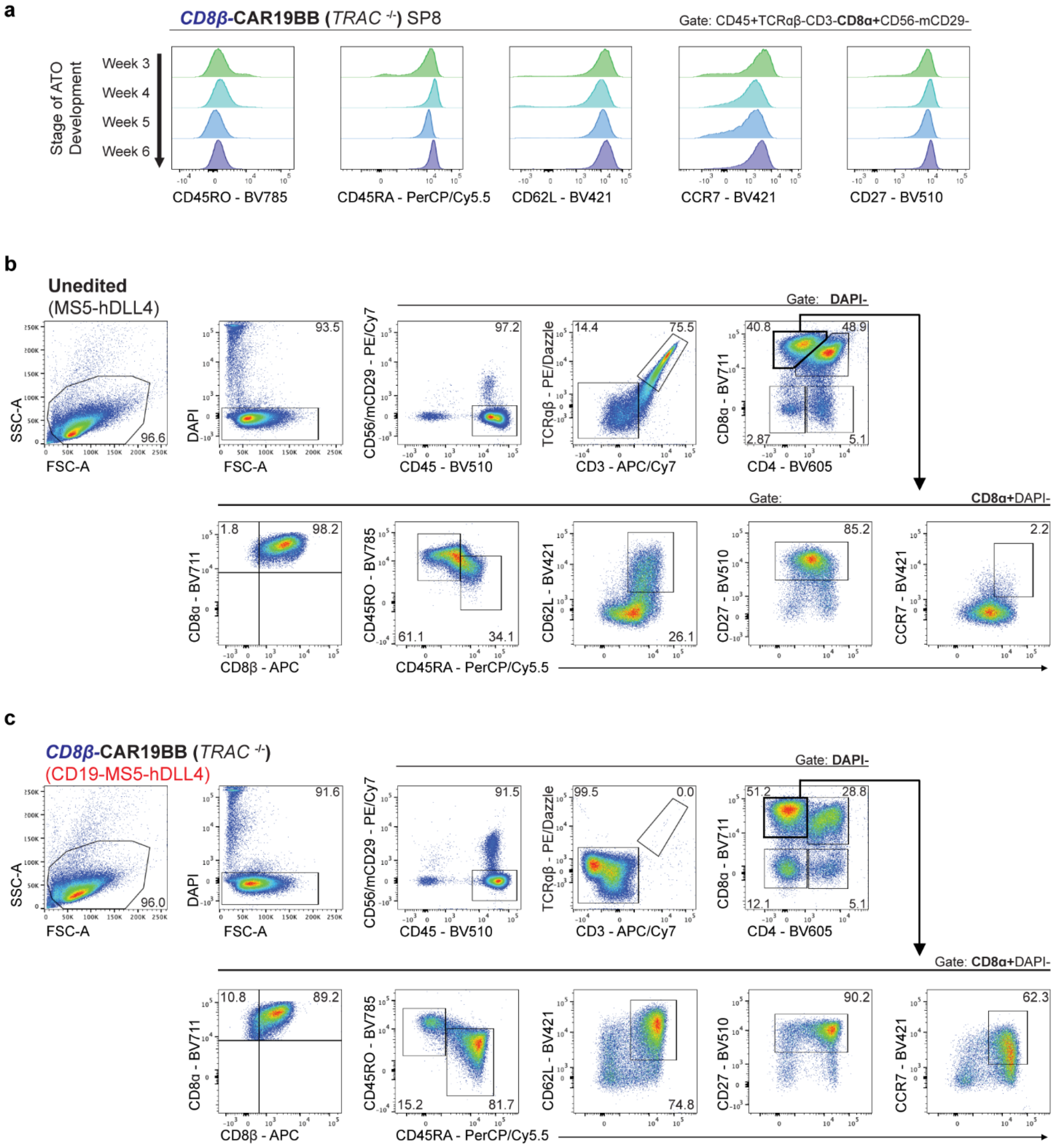


**Supplementary Figure 7: Development and phenotype of ATO cultures processed for single-cell sequencing**

**(a)** Phenotype of *CD8B*-CAR19BB (*TRAC*$^{-/-}$) SP8 T cells analyzed at weeks 3 through 6 of CD19-expressing ATO cultures (data representative of n = 4 independent experiments).

**(b,c)** Analysis of whole ATO cultures and phenotype of SP8 T cells processed for nuclei isolation and submitted for sequencing at week 3 of culture. **(b)** Unedited T-iPSCs were differentiated using MS5-hDLL4 and **(c)** *CD8B*-CAR19BB (*TRAC*$^{-/-}$) T-iPSCs were differentiated using CD19-MS5-hDLL4 (data representative of n = 2 independent experiments).

### Supplementary Figure 8: Single-cell, multi-modal analysis of unedited and *CD8B*-CAR19BB (*TRAC*$^{-/-}$) ATO cultures

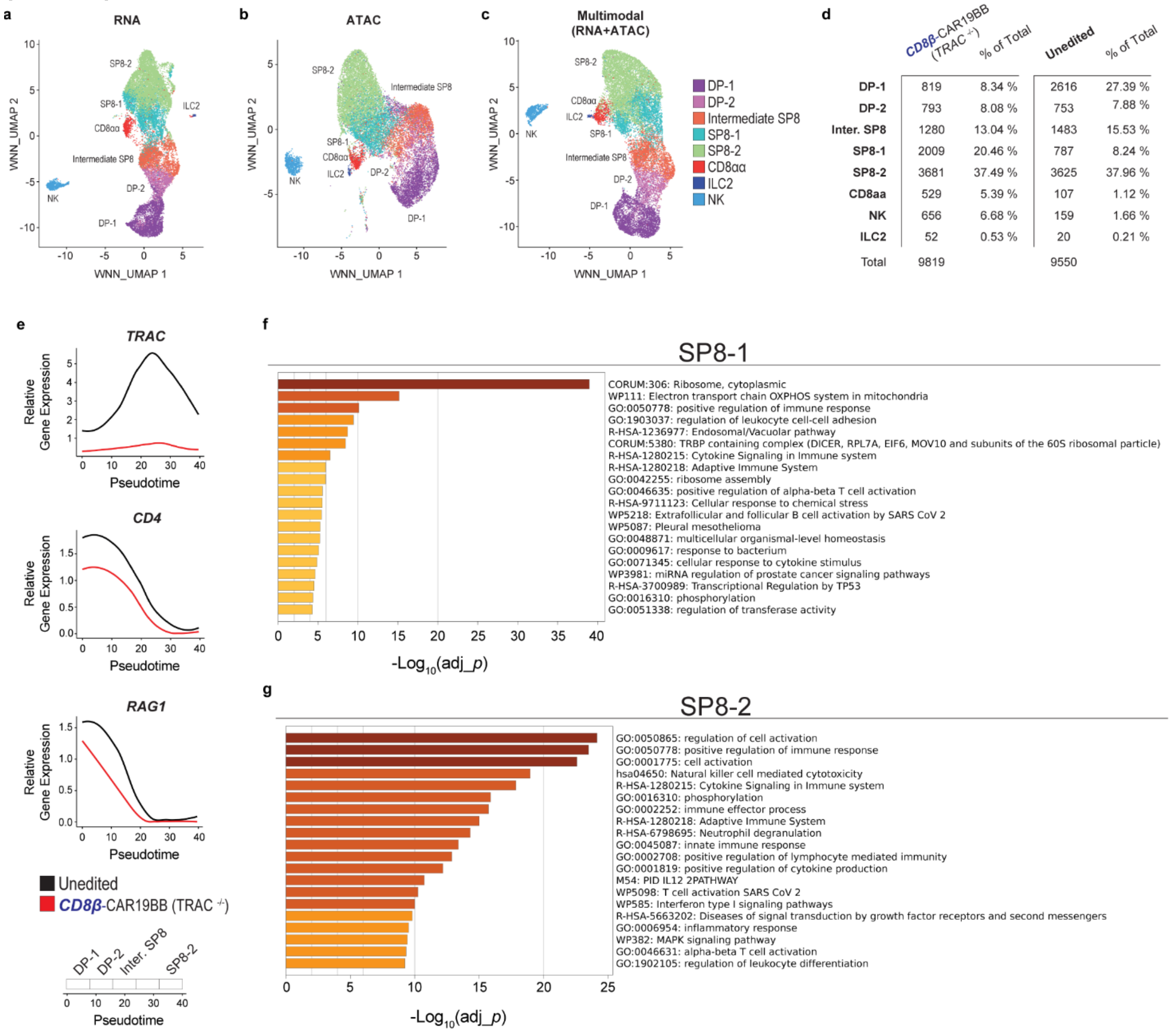


**Supplementary Figure 8: Single-cell, multi-modal analysis of unedited and *CD8B*-CAR19BB (*TRAC*$^{-/-}$) ATO cultures**

**(a-c)** Combined [unedited and *CD8B*-CAR19BB (*TRAC*$^{-/-}$)] WNN UMAP visualization of annotated populations generated using **(a)** RNA-seq only, **(b)** ATAC-seq only, and **(c)** Multimodal data (integration of RNA and ATAC libraries) sequenced at week 3 of ATO cultures.

**(d)** Number and proportion of cells captured during sequencing in each annotated population identified from unedited and *CD8B*-CAR19BB (*TRAC*$^{-/-}$) ATOs at week 3 of culture.

**(e)** Relative gene expression of *TRAC, CD4,* and *RAG1* across pseudotime in unedited (black) and *CD8B*-CAR19BB (*TRAC*$^{-/-}$) (red) T cells.

**(f,g)** Top 20 representative enriched terms across differentially expressed genes between **(f)** SP8-1 and **(g)** SP8-2 clusters. Each bar represents a –log10-transformed adjusted p-value. Gene enrichment analysis was performed using Metascape (https://metascape.org)[24].
